# Targeting immunometabolic pathways with AZD1656 alleviates inflammation and metabolic dysfunction in type 2 diabetic cardiomyopathy

**DOI:** 10.1101/2025.04.10.648128

**Authors:** Stephanie Anderson, Anja Karlstaedt, Megan Young, Loucia Karatzia, Fenn Cullen, Jianmin Chen, Caroline E. O’Riordan, Michael R. Barnes, Zorana Štaka, Lauren J. Albee, Conor Garrod-Ketchley, Sanushi Dambure, Hiran A. Prag, Filip Cvetko, Jack J.J.J. Miller, Christoph Thiemermann, Andrew JM. Lewis, Michael P. Murphy, David M. Smith, Sian M. Henson, Damian J. Tyler, Dunja Aksentijevic

**Affiliations:** Department of Physiology, Anatomy and Genetics, University of Oxford, Oxford, UK; Department of Cardiology, Smidt Heart Institute, Cedars-Sinai Medical Center, Los Angeles, USA; William Harvey Research Institute, Bart’s and The London Faculty of Medicine and Dentistry, Queen Mary University of London, London, UK; Faculty of Electrical Engineering, University of East Sarajevo, East Sarajevo, Bosnia and Herzegovina; School of Cardiovascular Science and Medicine, The Rayne Institute, St Thomas Hospital, King’s College London, London, UK; MRC Mitochondrial Biology Unit, University of Cambridge, Cambridge, UK; The MR Research Centre and The PET Centre, Clinical Medicine, Aarhus University, Aarhus 8200, Denmark; Clarendon Laboratory, Department of Physics, University of Oxford, UK; Radcliffe Department of Medicine, University of Oxford, OX3 9DU, UK; Emerging Innovation Unit, Discovery Sciences, R&D, AstraZeneca, Cambridge, UK

**Keywords:** type 2 diabetic cardiomyopathy, inflammation, metabolic remodelling, cardio-immunology, type 2 diabetes

## Abstract

Type 2 diabetes (T2D) precipitates diabetic cardiomyopathy (dbCM), a condition characterized by chronic inflammation, metabolic dysregulation and impaired cardiac performance. Here we show that the glucokinase activator AZD1656, originally developed for glycaemic control but later identified to have immunomodulatory effects, reverses cardiac dysfunction and metabolic remodelling in dbCM. In obese, hyperglycaemic db/db mice with diastolic dysfunction, six weeks of AZD1656 treatment improved myocardial performance, reduced infarct size and enhanced post-ischaemic recovery. Integrated metabolic, functional and histological analyses revealed restoration of mitochondrial metabolism and attenuation of fibrosis. Mechanistically, AZD1656 remodelled the cardiac immune landscape by promoting regulatory T-cell infiltration. These findings demonstrate a link between cardiac inflammation and metabolic remodelling in dbCM and highlight that modulation of immune cells and metabolism can protect the diabetic heart. Targeting immunometabolic pathways may therefore offer a therapeutic strategy to alleviate cardiac dysfunction and reduce infarct vulnerability in T2D

## Introduction

Diabetic cardiomyopathy (dbCM) is a complication of type II diabetes (T2D), characterized by impaired cardiac function, disrupted metabolism, systemic inflammation and, consequently, an increased incidence of ischemic heart disease and heart failure^1^. The dbCM heart is metabolically inflexible relying heavily on fatty acid (FA) oxidation. This loss of metabolic flexibility means the dbCM heart is chronically energetically impaired^2^. Whilst metabolic derangements are a well-established risk factor for a worsening prognosis in dbCM, currently there are no therapeutic agents specifically designed to target cardiac metabolism in dbCM.

Glucokinase and hexokinase catalyse glucose phosphorylation to glucose-6-phosphate in the first step of glycolysis. While hexokinase is found in most tissues, glucokinase is primarily found in the pancreatic β-cells, the liver, and a subset of T cells. In these tissues, glucokinase acts as a glucose sensor and regulates glucose-stimulated insulin secretion and hepatic glucose uptake. As such, activating glucokinase is an attractive target for the treatment of T2D. To this end, AZD1656 was developed as a specific activator of glucokinase with >100-fold selectivity over hexokinase and was expected to provide effective glycaemic control in T2D through its activity in pancreas and liver^3^. However, in twenty-five clinical trials involving 900 T2D patients^4^ AZD1656 failed to fulfil its primary endpoint of controlling hyperglycaemia, only improving glucose levels for up to 4 months.

In a separate study, AZD1656 was shown to exert immunoregulatory influences on metabolic pathways governing regulatory T-cell (Treg) motility by Kishore et al.^5^, who demonstrated that activation of glucokinase (GCK) enhanced Treg migration to inflamed tissues. In Tregs, engagement of the CD28 costimulatory receptor activates PI3K-mTORC2 signalling, which couples metabolic reprogramming with cytoskeletal remodeling^5^. Through Akt phosphorylation, this cascade promotes the transcription and activation of GCK and modulates its regulatory partner GCKR, releasing GCK to participate in metabolic and migratory processes. Freed from GCKR inhibition, GCK translocates toward the cell periphery, where it supports localised glycolytic ATP production essential for actin remodelling and cell motility^5^. Pharmacological activation of this pathway by AZD1656, a selective GCK activator, thus enhances Treg glycolytic capacity and motility, fostering an anti-inflammatory, tissue-protective immune phenotype. Importantly, GCK activation enhances Treg trafficking without affecting proliferation^5^, confirming that increased glycolytic flux, rather than cell division, underlies their migratory advantage.

Clinical data reinforce this immunometabolic mechanism. In the ARCADIA trial^6^ (NCT04516759), treatment with AZD1656 improved outcomes in patients with diabetes and systemic inflammation, reducing mortality and accelerating recovery. These benefits were accompanied by a shift toward immune homeostasis, including decreased circulating cardiotropic cMet□ Tregs, consistent with enhanced Treg migration into inflamed tissues^6^. Genetic evidence further supports this model with individuals carrying the GCKR P446L loss-of-function polymorphism, which diminishes inhibition of GCK, showing increased hepatic GCK activity, altered glucose and lipid metabolism, and enhanced Treg motility independent of glucose uptake or Akt signaling^5^. Together, these data identify GCK-dependent glycolysis as a critical determinant of Treg trafficking.

Building on this immunomodulatory foundation, the ADOPTION trial (NCT05216172), is a placebo-controlled, double-blind randomised study investigating AZD1656 as a novel strategy to modulate T-cell immunometabolism in renal transplant recipients with type 2 diabetes. The observed Treg-specific actions of AZD1656 formed the basis for its evaluation in the ADOPTION-CKD trial designed to test whether glucokinase activation can modulate systemic inflammation and improve outcomes in chronic kidney disease.

It is well known that chronic inflammation is a hallmark of T2D, accompanied by increased circulating levels of highly inflammatory senescent T-cells^1^. Proinflammatory stimuli such as systemic metabolic stress in T2D (hyperinsulinemia, hyperlipidaemia, hyperglycaemia, oxidative stress) may trigger T cell-mediated autoimmunity leading to chronic inflammation in the heart^7^. Despite this, many anti-inflammatory agents have failed to improve cardiac function in clinical trials as they mostly target innate immunity and acute inflammation^7,8^. Systemic hyperlipidaemia characteristic of T2D has also been shown to promote the differentiation of pro-inflammatory effector memory T-cells (Tem), while reducing the number of regulatory T-cells, essential for immune homeostasis^9^. In chronic metabolic stress states such as T2D, high circulating fatty acids have been shown to drive the differentiation of CD4+ Tem and migration into non-lymphoid tissues thus promoting inflammation by pro-inflammatory T-cell cytokine release and systemic inflammasome activation^7,9^. However, whether systemic T-cell-mediated inflammation in T2D leads to myocardial inflammation and whether it impacts metabolic derangement in dbCM remains unknown.

Given the potential immuno-modulatory effect of AZD1656, we set out to test the immune-regulatory potential of AZD1656 during cardiac metabolic and functional derangement in T2D. We show that a 6-week AZD1656 drug intervention in the db/db mouse model of dbCM attenuated diastolic dysfunction, reversed metabolic remodelling and reduced infarct size versus untreated dbCM. The beneficial cardiometabolic effects of AZD1656 treatment were not driven by systemic metabolic alterations in the liver, muscles or adipose tissue but AZD1656 did reduce cardiac inflammation and fibrosis.

AZD1656 also normalizes myocardial gene expression including those mediating inflammation as well as governing key pro-survival and metabolic pathways (i.e HIF-1α, Nrf2, sirtuin, PPARα, apoptosis). Consequently, AZD1656 treatment may represent a new therapeutic option for dbCM by targeting immunometabolism to reduce cardiac inflammation and improve cardiac remodelling.

## Materials and Methods

### Animals

Commercially available type 2 diabetic mice (db/db mouse, The Jackson Laboratory homozygous BKS.Cg-Dock7^m^ +/+ Lepr^db^/J, male, Charles River, Italy) were purchased at 8 weeks with corresponding lean littermate controls (heterozygous Dock7^m^ +/+ Lepr^db^, Charles River, Italy). Animals were kept under pathogen-free conditions, 12h light–dark cycle, controlled temperature (20–22°C), and fed chow and water *ad libitum*. Circulating glucose (tail sample, Accu-Check, Roche) and body weight were monitored weekly. This investigation conformed to UK Home Office Guidance on the Operation of the Animals (Scientific Procedures) Act, 1986.

### AZD1656 drug treatment protocol

AZD1656 (AZD, Astra Zeneca, UK) is a selective glucokinase activator. No significant clinical effects nor specific toxicology risks other than hypoglycaemia were noted in clinical trials^10^. The db/db and lean controls were divided into 3 groups at 13 weeks of age: Group I - lean controls, Envigo diets control diet (2019 Teklad Global 19% Protein Rodent Diet, irradiated, Teklad Custom Diets, Envigo), Group II - db/db, Envigo diets control diet and Group III - db/db, AZD1656 diet (30 mg/kg body weight/day, Envigo Specialist Diets, USA; drug dosing based on^11,12^), Figure 1a. Diets were fully matched in terms of nutritional standardisation and food intake was not affected by dietary drug incorporation. Animals (study total: Group I - Controls n=64, Group II - db/db n=63, Group III – AZD1656 n=57) were kept on the feeding protocol from 14 until 20 weeks of age when the study reached endpoint. For the obesity induction protocol, C57/BL6 8-week male mice (n=18, Charles River, UK) were fed a high-fat diet for 12 weeks *ad libitum* (HFD, cat. no. TD.200185; Envigo, 60.3% fat, 21.4% carbohydrate, and 18.3% protein; percentage of energy in kcal).^13^ *In vivo* physiology and heart function as well as *ex vivo* cardiac metabolome were assessed as described before^13^.

**Figure 1.**
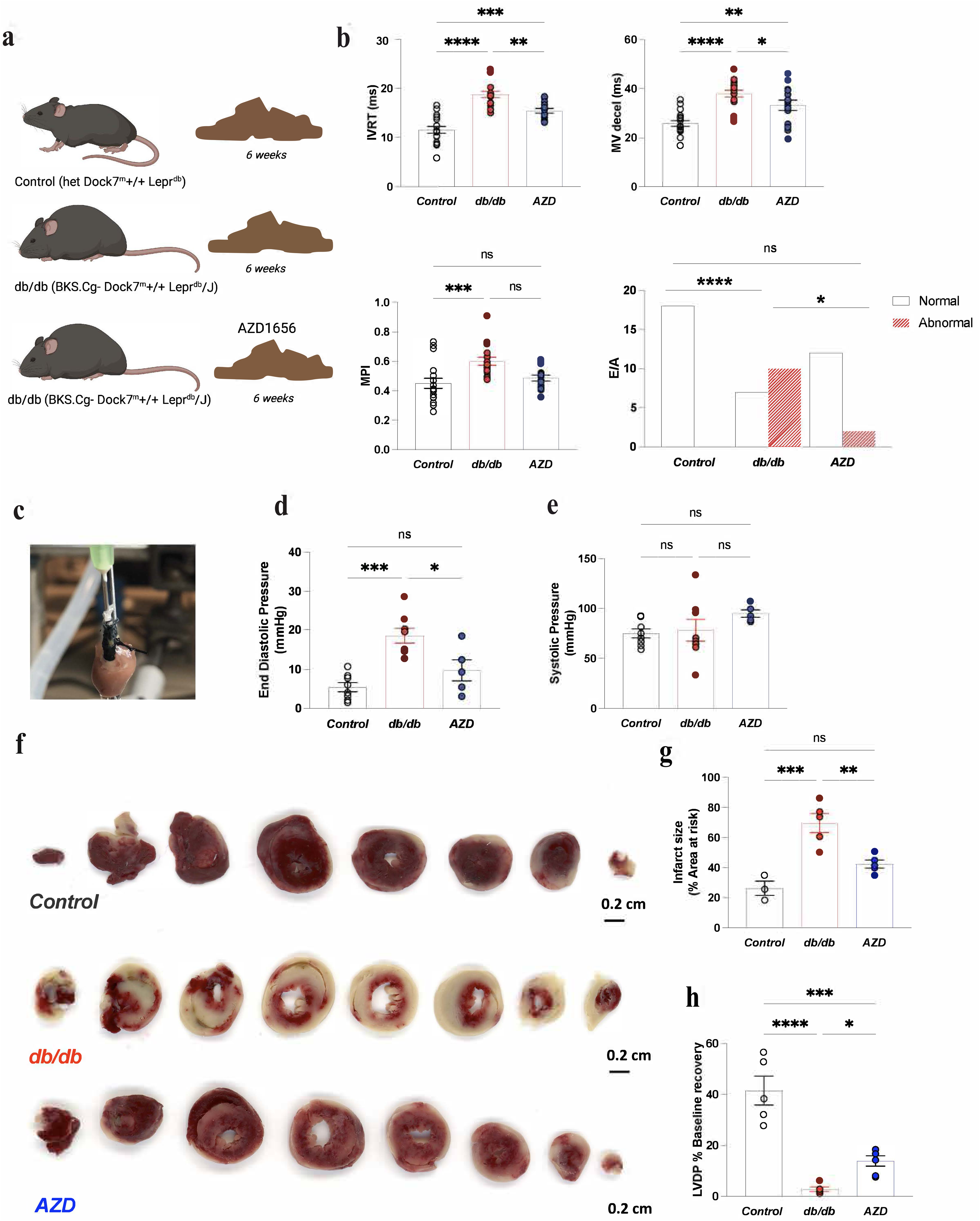
AZD1656 treatment attenuates diastolic dysfunction and reduces infarct size in dbCM. **a**) Graphical summary of the AZD1656 treatment protocol. Mice were fed nutritionally standardised and matched Teklad Standard Base Diet (Envigo) with the addition of AZD1656 to the diet used for db/db treatment group (30mg/kg) **b**) *In vivo* echocardiography assessment of diastolic dysfunction: IVRT, MV decel, MPI, mitral valve E/A control vs db/db P<0.0003, AZD vs control P<0.185 AZD vs db/db P<0.036 by Fischer’s exact test. Statistical analysis approach for E/A analysis described in Methods. (control n=17, db/db n=17, AZD n=14) **c**) representative Langendorff perfused heart **d**) **e**) Langendorff perfused heart function data (Control n=9, db/db n=9, AZD n=6) **f**) Representative TTC stained cardiac cross sections post 20 min total global ischemia and 2 hr reperfusion used for quantification of the infarct size. **g**) Quantification of infarct size post I/R injury (db/db, AZD n=5/group, control n=3) **h**) Improved LVDP recovery post ischemia in Langendorff perfused hearts (Control n=5, db/db n=5, AZD n=6) f) *P<0.05, ***P<0.01 by ANOVA

### Body Composition analysis

Body composition analysis was carried out by non-invasive magnetic resonance relaxometry using an EchoMRI^TM^ Body Composition Analyser E26-348-MT (Houston, Texas, USA). Accumulation factor was for extra-high precision (3×) resulting in a scan time of approximately 2.5 min.

### In vivo Assessment of left ventricular systolic and diastolic function

#### CINE Magnetic Resonance Imaging

For the assessment of systolic function, mice were imaged in a 7T MRI instrument (Agilent, USA) using CINE magnetic resonance imaging, as previously described^14^. Eight to 10 short-axis slices (slice thickness, 1.0 mm; matrix size, 256 x 256; field of view, 25.6 x 25.6 mm; echo time/repetition time, 0.3/4.6 ms; flip angle, 30°; and number of averages, 4) were acquired with a gradient echo, fast low-angle shot sequence^15^. Left ventricular volumes were derived using the freehand draw function in ImageJ (National Institutes of Health). For each heart, left ventricular mass, ejection fraction, stroke volume, and cardiac output were calculated.

#### Echocardiography

M-mode and Doppler echocardiography was performed in 20-week lean controls, db/db and AZD1656 treated db/db mice at the end of the drug treatment protocol as previously described^13,16,17^. Anaesthesia was induced with 4% isoflurane and maintained at 1.5-2% for the duration of the procedure. Before assessment of cardiac function, fur was removed from the chest area to allow accurate assessment of cardiac function and mice were allowed to stabilize for at least 10 minutes. Body temperature was maintained at 37°C. During echocardiography, heart rate was measured from electrocardiogram. Echocardiography images were recorded using a Vevo-3100 imaging system with a 40-MHz linear probe (VisualSonics, Toronto, Canada). Morphological measurements were taken in the parasternal short axis view at the level of the papillary muscles and ejection fraction was calculated from M-mode images. Diastolic transmitral left ventricle (LV) inflow images were acquired from apical four-chamber views using pulsed-wave doppler to calculate early (E) and late (atrial, A) peak filling blood flow velocities and E-wave deceleration time. Analysis was performed using VevoLab 5.5.1 software. Cardiac function was assessed blinded to the phenotype.

#### Hyperpolarized Magnetic Resonance Spectroscopy

For the *in vivo* assessment of cardiac metabolism, hyperpolarized magnetic resonance spectroscopy was used to monitor the down-stream fate of hyperpolarized [1-^13^C]pyruvate as previously described^18^. Experiments were performed between 7 and 11 A.M. when mice were in the fed state. A total of 40 mg [1-^13^C]pyruvic acid (Sigma-Aldrich) doped with 15 mmol/L trityl radical (OXO63; GE Healthcare) and 3 mL Dotarem (1:50 dilution; Guerbet) was hyperpolarized in a prototype polarizer, with 30-60 min of microwave irradiation^19^. The sample was subsequently dissolved in a pressurized and heated alkaline solution, containing 2.4 g/L sodium hydroxide and 100 mg/L EDTA dipotassium salt (Sigma-Aldrich), to yield a solution of 80 mmol/L hyperpolarized sodium [1-^13^C] pyruvate with a polarization of ∼30%. A total of 200 μL was injected over 10 s via the tail vein. ^13^C MR pulse-acquire cardiac spectra were acquired over 60 s following injection of hyperpolarized [1-^13^C]pyruvate (repetition time, 1 s; excitation flip angle, 15°; sweep width, 13,021 Hz; acquired points, 2,048; and frequency centred on the C1 pyruvate resonance)^18^. The ^13^C label from pyruvate and its metabolic products was summed over 60 s from the first appearance of pyruvate and fitted with the AMARES algorithm in jMRUI^20^. Data are reported as the ratio of metabolic product to the [1-^13^C]pyruvate signal to normalize for differences in polarization and delivery time.

#### Langendorff heart perfusions

Mice were terminally anesthetized, hearts rapidly excised, cannulated and perfused as a standard Langendorff preparation as previously described^21^. Krebs-Henseleit (KH) perfusion buffer was continuously gassed with 95% O2/5% CO2 (pH 7.4, 37 °C) containing (in mM): NaCl (116), KCl (4.7), MgSO4.7H2O, KH2PO4 (x), (1.2), NaHCO3 (25), CaCl2 (1.4), and enriched with metabolites [glucose (11), intralipid 0.4mM, 1 sodium L-lactate; 0.1 sodium pyruvate; 0.5 L-glutamic acid monosodium salt monohydrate; 5□mU□l^−1^insulin (NovoRapid insulin, Novo Nordisk, Denmark),] and paced at 550bpm via epicardial silver wire electrodes placed at the apex of the left ventricle and the right atrium. The impact of 1μM AZD^5^ on heart function and metabolism was assessed in unpaced hearts perfused using hyperglycaemic (11mM) crystalloid KH buffer prepared in amber glassware immediately prior to the experiment. At the end of each experiment, hearts, liver, adipose tissue and muscle (gastrocnemius and soleus) were immediately freeze-clamped using Wollenberger tongs for metabolic profiling.

Myocardial infarct size was quantified as described before^22^. In brief, after 20 min of equilibration, Langendorff crystalloid KH buffer perfused hearts were subject to 20 min of global normothermic ischemia (37°C) and 2 hours of reperfusion (37°C). At the end of the protocol, hearts were perfused for 10 mins with 3% triphenyltetrazolium chloride (TTC) in KH Buffer followed by 10 min incubation in 3% TTC-KH. Tissue was sectioned (mouse heart gauge, Zivic instruments, USA) and infarct field was quantified using ImageJ Software (National Institute of Health).

#### Metabolomic profiling

Snap frozen and pulverized tissue [heart, liver, skeletal muscle(gastrocnemius+soleus), adipose tissue] was analysed using ^1^H NMR high resolution spectroscopy as previously described^23,24^. Non-targeted lipid profiling of the cardiac tissue was performed by LC-MS MS by School of Chemical and Physical Sciences Mass Spectrometry Laboratory Services. Total lipids were extracted from a weighed amount of heart tissue using a modified Folch extraction protocol. Briefly, each tissue sample was homogenized by probe sonication in 3.75 mL of chloroform:methanol (1:2, v/v). Subsequently, 1.25 mL of chloroform was added, and the organic phase was separated from tissue debris following additional probe sonication. The mixture was then washed with 2.25 mL of 0.9% NaCl solution to induce phase separation. The lower organic phase containing the extracted lipids was collected by centrifugation and evaporated to dryness under a gentle stream of nitrogen. The resulting lipid extract was reconstituted in 100 μL of methanol and subjected to HPLC-MS analysis. Total lipids were separated using an ACQUITY UPLC system (Waters Corporation, USA) equipped with a UPLC CSH C18 column (2.1 × 100 mm, 1.7 μm (Waters Corporation, USA), maintained at 55 °C. The mobile phases consisted of (A) acetonitrile:water (60:40, v/v) containing 10 mM ammonium formate and 0.1% formic acid, and (B) isopropanol:acetonitrile (90:10, v/v) containing 10 mM ammonium formate and 0.1% formic acid. Lipid species were separated using a step gradient to 70% solvent B over 12 min at a flow rate of 0.45 mL/min.

Mass spectrometric detection was performed on a Synapt G2-Si high-resolution mass spectrometer (Waters Corporation, USA) operated in positive electrospray ionization (ESI) mode. Mass spectra were acquired over an *m/z* range of 50–1200. Nitrogen was used as both the cone and desolvation gas, with flow rates of 25 L/h and 900 L/h, respectively. The source and desolvation temperatures were maintained at 125 °C and 350 °C. Leucine enkephalin (Leu-Enk) was continuously infused and used as a lock mass for real-time mass calibration. Data acquisition and instrument control were performed using MassLynx software (Waters Corporation,USA). Cardiotonic lipids were analysed in a non-targeted acquisition mode. Following mass spectrometry data collection, samples were processed using UNIFI software (Waters Corporation, USA). The observed mass intensities of each identified lipid were matched against a curated cardiolipid library comprising approximately 270 lipid species. Lipid signal intensities were normalized to the weight of the corresponding heart tissue samples. Annotation of lipid species to lipid classes and categories was conducted using Lipidmaps^25^. MS intensities were then used to weight the molecular fatty acyl (FA) contributions and calculate class wise and total FA profiles. The carbon chain length distribution of phospholipid species was expressed as a percentage of the total signal within each class [carbon length (%)]. For each phosphatidylcholine (PC) species, the total number of carbons across both fatty acyl chains was calculated, and the proportion of all PC species containing that total carbon number was determined. Similarly, the degree of unsaturation was assessed by calculating the total number of double bonds in the fatty acyl chains of each PC species and expressing the proportion of species carrying that total number of unsaturations as a percentage of all PC species [carbon bonds (%)].

#### Lipoprotein lipase assay

Lipoprotein lipase (LPL) activity was measured in cardiac tissue samples (control, db/db n=10/group, AZD n=9) using a fluorometric assay according to the manufacturer’s protocol (Cell Biolabs, STA-610, Lot:8724005). Briefly, cardiac tissue samples were homogenised in 20mM Tris (pH 7.5), 150mM NaCl before centrifugation at 10,000 × g for 10 minutes at 4°C. The supernatant was diluted 1:50 in LPL assay buffer and run alongside an LPL enzyme standard curve. Fluorescence intensities were measured at 485 excitation and 520 emission.

#### In silico analysis of cardiac metabolism

In silico simulations were performed using the metabolic network of the cardiomyocyte, CardioNet^21^. Mathematical modelling has previously been used to study the dynamics of cardiac metabolism in response to stress, and CardioNet has been successfully applied to identify limiting metabolic processes and estimate flux distributions^21,26,27^. Optimization problems were defined with the objective to minimize the total sum of fluxes through the CardioNet metabolic network^28^. Simulations for control, db/db or AZD1656 groups were based on the assumption that cardiomyocytes seek to maintain a certain ATP provision to sustain cardiac contractile function alongside the synthesis of macromolecules, including structural proteins and phospholipids for membranes^28^. We included the intracellular metabolite concentrations of 23 metabolites: NADH, formate, ATP/ADP pool, fumarate, glucose, (phospho)creatinine, glycine, taurine, carnitine, (phospho)choline, acetyl-carnitine, aspartate, glutamine, glutamate, succinate, acetate, alanine, lactate, valine and (iso)leucine. Furthermore, we constrained extracellular metabolites that could be taken up from the blood, including glucose, lactate, cholesterol and free fatty acids. Linear programming was solved using the GUROBI solver (version 9.1.2 build v9.1.2rc0, Linux64)^29^.

Details of all reactions and their metabolic subsystems used for CardioNet analysis were as classified in the Kyoto Encyclopedia of Genes and Genomes database.^30^

#### Tissue extraction and digestion for fluorescence-activated cell sorting

To generate leukocyte single cell suspensions for characterisation of immune cell populations, hearts were isolated from mice and digested. Mice were euthanised using an overdose of anaesthesia with 5 % isoflurane in 2 L/min O_2_. Cessation of pedal and corneal reflexes were checked, and death was confirmed by cervical dislocation.

Cardiac tissue suspensions were prepared by perfusing hearts with cold HBSS for 5 min prior to removing the atria and mincing the ventricular tissue that was digested in collagenase I (Worthington Laboratories, C1639, 450 units/mL), collagenase XI (Worthington Laboratories, C7657, 125 units/mL), DNase 1 (Worthington Laboratories, D4527, 60 units/mL) Hyaluronidase (Sigma Aldrich, H3506, 60 units/mL) and 20mM Hepes in PBS for 20 minutes at 37°C with gentle agitation (Thermomixer, 750 rpm). Spleens were mechanically dissociated by mashing through a cell strainer using the plunger end of a sterile syringe. Samples were passed through a 70 µm cell strainer, rinsed with 10 mL cold 2% FBS/ PBS and centrifuged at 400g for 10 min at 4°C. Supernatant was removed and pellets resuspended in 5 mL red blood cell lysis buffer (BioLegend) and incubated on ice for 10 min with occasional agitation after which 10 mL cold 2% FBS/ PBS was added to neutralise the lysis. Samples were then centrifuged for 8 min at 4°C and 400g, supernatant was removed and cells were resuspended and incubated with FC-block (BioLegend 101320, 1 µL per 1 × 10^6^/ mL cells). Cells were then washed again and resuspended in PBS ready for counting and antibody staining.

#### Flow cytometry

Cells isolated from tissues were resuspended (∼10^7^/ml) and incubated for 30 minutes at room temperature with fluorochrome-conjugated antibodies (Supplementary Table 1) in 100 μl of flow cytometry buffer made of PBS containing 0.1% sodium azide (SigmaAldrich) and 1% FBS. For intracellular marker staining, cells were fixed and permeabilized for 30 minutes at 4°C using fixation/permeabilization kit (eBioscience), washed in 1X permeabilization buffer (eBioscience) and stained with fluorochrome-conjugated antibodies in 1x permeabilization buffer for 30 minutes at 4°C. A final wash with 1x permeabilization buffer was performed, centrifuged and resuspended in 200μl of flow cytometry buffer. Alternatively, cells were fixed (Fix/Perm kit, BioLegend 426803) and stored at 4°C. Cell viability was assessed using incubation with viability dyes (Supplementary Table 1). Samples were analysed on FACSAriaIII (BD Biosciences) running FACSDiVa v.8.0 software (BD Biosciences). CD3 beads (Miltenyi, UK) were routinely used to calibrate the cytometer. Single stain and fluorescence minus one control were acquired for compensation and precise gating (Supplementary Fig1-4 gating strategies). Compensation was automatically calculated, and samples analyzed using FlowJo software (version 10, FlowJo LLC, Oregon, USA).

#### Histology

Hearts were collected into buffered formalin (Sigma Aldrich). All histological processing was carried out by the Bart’s Cancer Institute Histology Core Facility. For the assessment of cardiac fibrosis, paraffin embedded sections were stained using Masson’s trichrome staining. Stained cardiac sections were scanned using a Nanozoom panoramic scanner (40x). Images (whole heart sections) were analysed using ImageJ (National Institute of Health).

### Electron microscopy

Left ventricle tissue samples were collected from db/db, AZD1656 treated db/db and lean control mice (n=3/group) and placed into phosphate buffered glutaraldehyde (4%) for fixing^26^. Samples were embedded in Araldite resin (Agar Scientific Ltd., Essex, UK) and ultrathin sectioned (Reichert-Jung Ultracut E Ultramicrotome, Leica, Vienna, Austria) for TEM analysis in JEM1400F (JEOL (UK) Ltd, Hertfordshire, UK) at 120 kV by Transmission Electron Microscope Facility, School of Biological and Behavioural Sciences. Image analysis was completed using Fiji Image J software, mitochondrial cristae surface density was calculated based on previously described methods^31^ ^32^.

### Plasma analysis

Blood samples were collected at the time of experimental endpoint into heparinized tubes. Plasma biochemical profiling was carried out by the MRC Mouse Biochemistry Laboratory (Addenbrookes NHS Hospital, Cambridge). Circulating plasma cytokines were assessed using XXL mouse cytokine array kit (Biotechne, USA) as described before^33^.

### RNA sequencing and Bioinformatic analysis

RNA extracted from the snap-frozen hearts was analysed by massive analysis of cDNA End (MACE-Seq). Rapid MACE-seq was used to prepare 3’ RNA sequencing libraries. Samples of 100 ng of DNA-depleted RNA were used for library preparation, using the Rapid MACE-Seq kit (GenXPro GmbH, Germany). Fragmented RNA underwent reverse transcription using barcoded oligo(dT) primers containing TrueQuant unique molecular identifiers, followed by template switching. PCR amplified libraries were purified by solid phase reversible immobilization beads and subsequent sequencing was performed using the Illumina platform NextSeq 500. Unprocessed sequencing reads were adapter-trimmed and quality-trimmed using Cutadapt (version 3.4,^34^). Deduplication based on UMIs (Unique Molecular Identifier) was performed using in-house tools. FastQC (0.11.9, ^35^) was used to assess the quality of sequencing reads. MultiQC (version 1.16,^36^) was used to create a single report visualising output from multiple tools across many samples, enabling global trends and biases to be quickly identified. MACE-Seq was annotated, reads quantified and p values for differential expression generated by GenXPro (Frankfurt, Germany).

Visual representations of the differentially expressed gene (DEG) dataset were performed using the Python programming language (version 3. 9.7) as well as libraries and packages, including: Matplotlib (version 3.4.3) for data visualization, Pandas (version 1.3.4) for data management, NumPy (version 1.20.3) for numerical computations, and Jupyter Notebook (version 6.4.5) for interactive code development. The functional enrichment analysis was performed using g:Profiler (version e108_eg55_p17_0254fbf) with g:SCS multiple testing correction method applying significance threshold of 0.05^37^. To determine enriched pathways and ontologies in all analysis comparisons, both Ingenuity Pathway Analysis (IPA; Ingenuity® Inc, Redwood city, CA) and Metascape comparison analysis (https://metascape.org)^38^ was performed on all genes with p<0.05. Comparisons were db/db_vs control (2740 DE genes) and AZ_vs_db/db (1271 DE genes). Both metascape and IPA utilise hypergeometric tests and Benjamini-Hochberg p-value correction to identify all ontology and pathway terms that contain a greater number of genes in common with an input list than expected by chance^38^.

### Data analysis and statistics

Data are presented as mean ± SEM. Comparison between groups were performed by Student’s t-test (Gaussian data distribution), two-way analysis of variance (ANOVA) with Bonferroni’s correction for multiple comparison and one-way ANOVA using Bonferroni’s correction for multiple comparisons where applicable. Normality of data distribution was examined using Shapiro–Wilk’s normality test. Statistical analysis was performed using GraphPad Prism (v9) software. Data analysis and visualization was conducted using R Studio (version 2022.12.0 Build 353). Partial Least-squares Discriminant Analysis (PLS-DA) was conducted using R package *mdatools*. Principal Component Analysis (PCA) was conducted with normalized intensities lipidomics data using the PCA function included in the *factoextra* package.

Flux estimations (CardioNet) were compared between experimental groups using Wilcoxon rank sum test and adjusted p-values were computed using the Bonferroni correction. Differences were considered significant when P < 0.05.

To assess the E/A ratio across groups, we employed a quantization approach due to the observed nonlinear (“U-shaped”) association between low and high E/A values with adverse outcomes, which is inconsistent with a linear relationship. Specifically, both low and high E/A values are known to be indicative of diastolic dysfunction^39^, which suggests a threshold effect (in which highly elevated E/A in the presence of disease is sometimes termed ‘pseudonormal’)^40^. We therefore discretised the observed E/A data into distinct categories, designating values more than three sample standard deviations above or below the mean of the lean control group as ‘abnormal’ (A), while those within the range of ±3 sample standard deviations were classified as ‘normal’ (N). This classification was chosen to reflect the nonlinear nature of the response, where extreme values on either end of the spectrum were associated with pathological conditions and is arguably more appropriate than a simple shift in means (as could be quantified e.g. via linear modelling and ANOVA type approaches), which would not adequately capture the complexities of the relationship. We then employed Fisher’s exact test for the resulting contingency table, which is particularly suited for categorical data comparisons in cases with small or unbalanced groups, implemented in the ‘RVAideMemoir’ R package^41,42^. Multiplicity correction was performed by the false discovery rate correction method of Benjamini & Hochberg^43^, with corrected p-values <0.05 considered significant.

## Results

### AZD1656 treatment improves diastolic function in the diabetic heart and offers protection from IR injury

The impact of 6-weeks of AZD1656 treatment (Figure 1a) on cardiac function in dbCM was assessed *in vivo* by CINE MRI and echocardiography (Table 1). As has previously been noted^44^, no reduction in systolic function was observed in untreated db/db mice, with a small but significant alteration in cine MRI-assessed ejection fraction observed alongside an unaltered cardiac output (Table 1). There was no evidence of pulmonary congestion or any increase in overall heart weight in db/db animals (Table 1). Cine MRI assessment (Table 1) identified unaltered end diastolic volume and decreased end systolic volume. Untreated db/db mice exhibited diastolic dysfunction typical for dbCM, evidenced by significant alterations in IVRT, MV deceleration time and E/A (Figure 1b, Table 1). *Ex vivo* functional assessment by Langendorff perfusion (Figure 1d) further confirmed diastolic dysfunction in db/db mice.

**Table 1.**
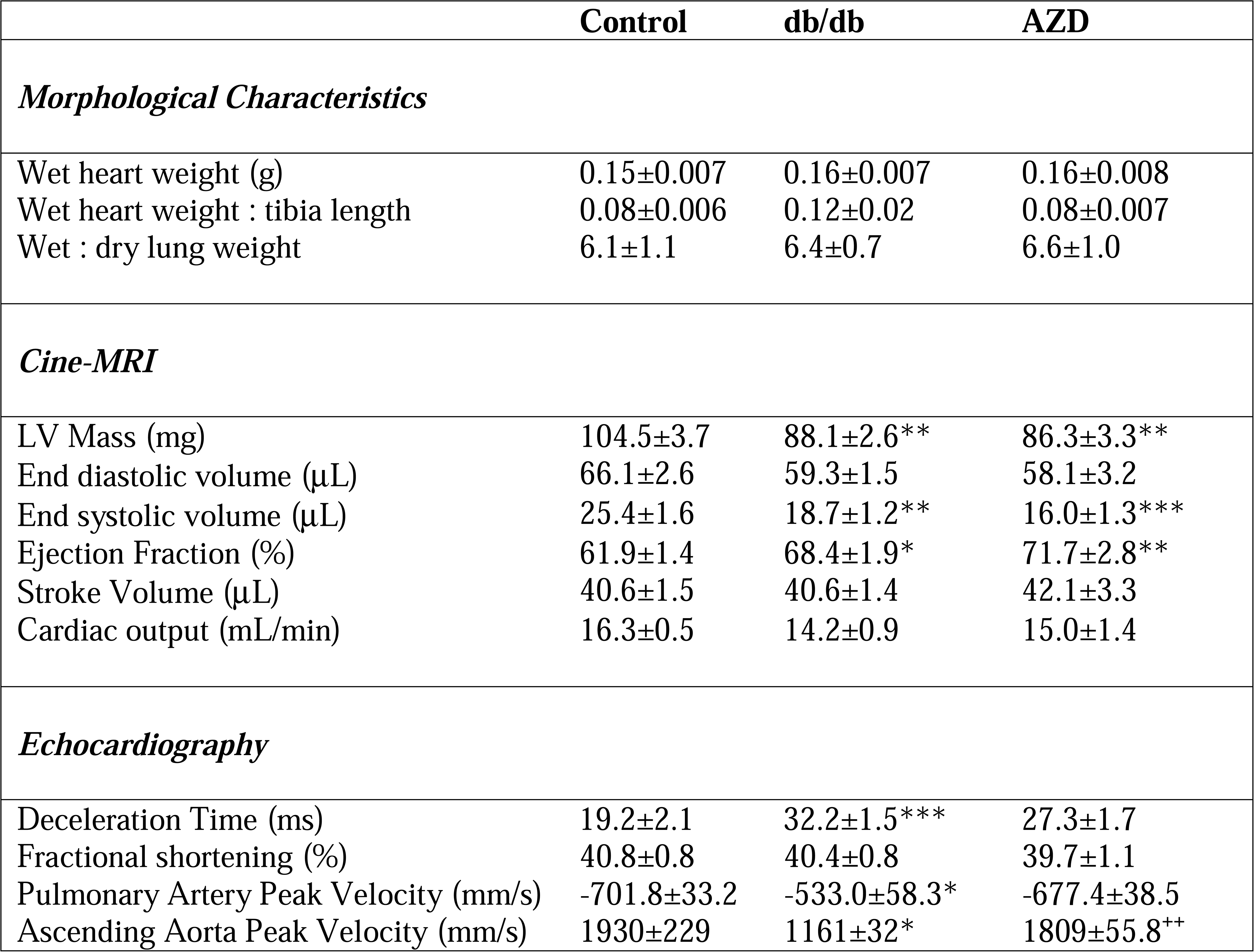
Morphological characteristics of the experimental model and impact of AZD treatment on in-vivo function assessed by Cine MRI and Echocardiography. Morphological Characteristics: wet heart weight (Control n=15, db/db n=15, AZD n=11), wet heart weight:tibia length (Control n=12, db/db n=10, AZD n=10).Cine MRI data (Control n=18, db/db n=15, AZD n=11). Echocardiography: deceleration time (n=5/group), Fractional shortening (Control n=10, db/db n=9, AZD n=10); Pulmonary Artery Peak Velocity (Control n=5, db/db n=3, AZD n=5), Ascending Aorta Peak Velocity (Control n=4, db/db n=3, AZD n=4), Kolmogorov-Smirnov test data normality test, multiple group comparison by one-way ANOVA with Bonferroni’s multiple comparison test. P<0.05, individual P values indicating level of significance stated in the table column (P value). ** P<0.005 db/db vs control, ***P<0.0005 db/db vs control, ^++^ P<0.05 AZD vs db/db

Treatment with AZD1656 enhanced *in vivo* cardiac function as improvements were noted in diastolic function, myocardial performance index (Table 1 Figure 1b), the doppler flow-assessed pulmonary artery VTI, pulmonary artery peak velocity (Table 1) and ascending aorta peak velocity (Table 1). This attenuation of myocardial dysfunction in db/db mice was accompanied by a protection from ischemia reperfusion injury as evidenced by reduced infarct size (Figure 1f, g) and significantly improved functional recovery post-myocardial infarction vs untreated db/db mice (Figure 1h).

### AZD1656 treatment improves cardiac metabolism in dbCM

Untreated db/db mice were characterized by severe cardiac metabolic dysfunction (Figure 2). Principal component analysis of the metabolomic profile, assessed by high-resolution ^1^H NMR spectroscopy (Figure 2a) confirmed that T2D leads to a distinct metabolomic profile of db/db hearts versus controls. Specifically, the metabolomic profile of T2D mice is characterized by a significant depletion of amino acids (glutamine, valine, glycine, taurine) and acetate, as well as altered TCA cycle intermediates (depleted fumarate, elevated succinate, Figure 2b). Collectively, metabolomic changes in db/db hearts were associated with reduction in energy reserves (phosphocreatine, ATP+ADP, PCr/ATP+ADP, Figure 2b).

**Figure 2.**
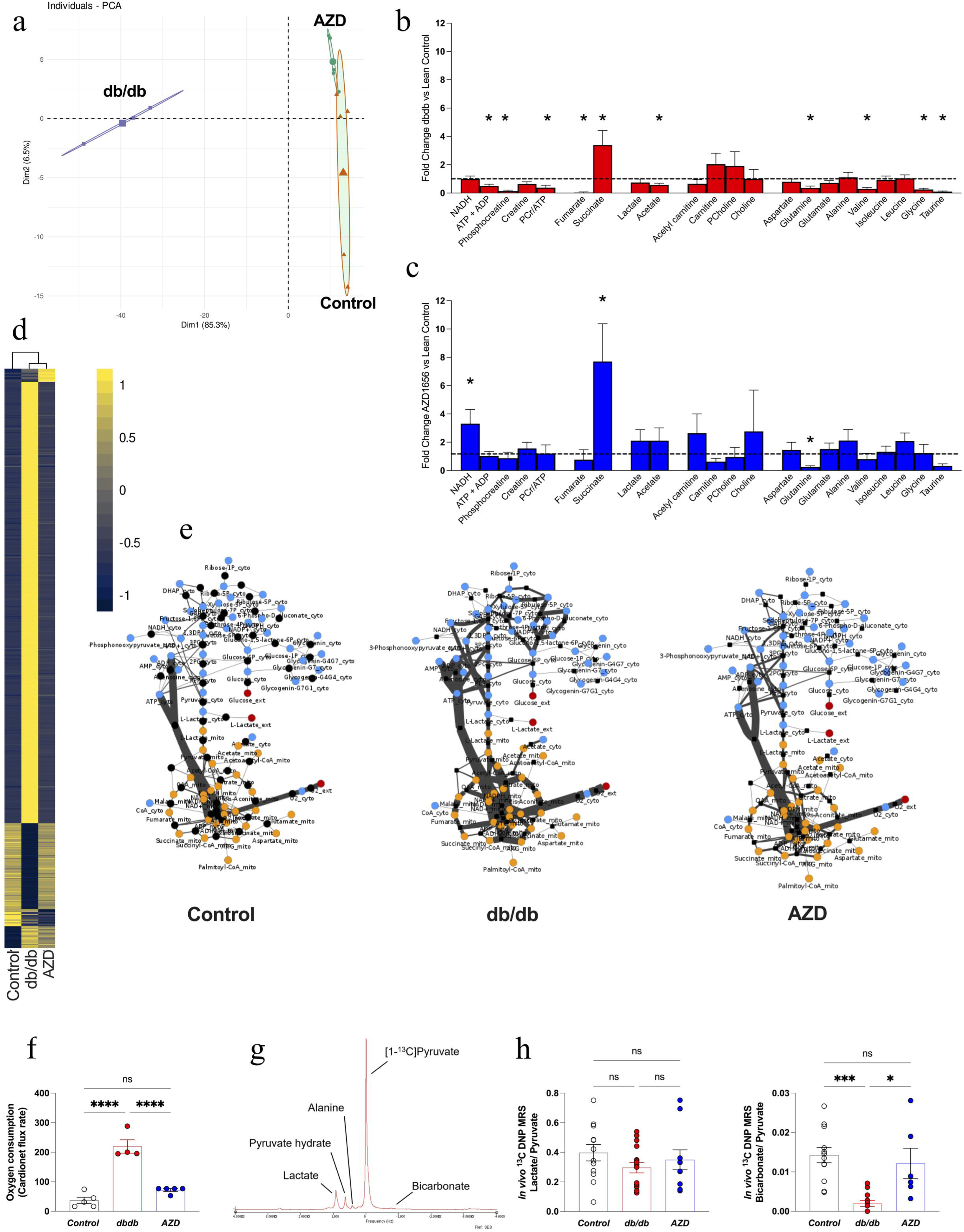
AZD1656 treatment improves cardiac metabolism in dbCM. **a**) PCA plot of ^1^H NMRs metabolomic profiling **b**) Fold change db/db vs control ^1^H NMR metabolomic spectroscopy profiling, n=6/group **c**) Fold change AZD treated db/db vs control ^1^H NMR metabolomic spectroscopy profiling, n=6/group **d**) Cardionet metabolic flux balance analysis based on the ^1^H NMR metabolomic profiling and plasma metabolomic analysis (n=6/group) **e**) Visualization of CardioNet metabolic flux predictions using Cytoscape. Metabolites and reactions are depicted as nodes and lines, respectively. The line thinckness corresponds with the calculate flux rate. **f**) Prediction of oxygen consumption rates based on CardioNet simulations. **g**) Representative annotated spectrum from *In vivo* ^13^C DNP MRS cardiac metabolic flux assessment **h**) *In vivo* ^13^C DNP MRS measured TCA cycle flux (Control n=12, db/db n=16, AZD n=10). Multiple group comparison by ANOVA. Two-tail comparison by student t-test. *P<0.05 ***P<0.01 ****P<0.001

In order to further analyse the cardiac metabolic profile in db/db mice, we applied a systems biology approach that combines metabolomics with constrained-based *in silico* modelling.^27,45^ To infer flux distributions, we integrated targeted metabolomics data from control, db/db and AZD1656-treated groups into flux balance analysis (FBA). Metabolic flux distributions were calculated using the mammalian network of cardiac metabolism, CardioNet^46^ based on quantified fold changes of metabolite concentrations between control and experimental groups as well as circulating metabolic profile. CardioNet *in silico* flux balance analysis of db/db cardiac metabolism further confirmed a markedly different metabolic profile compared to controls (Figure 2d). Computational simulations of db/db mice revealed enhanced anaerobic glycolysis and glucose flux through the pentose phosphate pathway, while glucose oxidation in the Krebs cycle was decreased (Figure 2e). The shift in extracellular nutrient supplies enhanced lipid oxidation (Figure 2e), which increased reactive oxygen species generation (Figure 2e) and markedly elevated oxygen consumption (Figure 2f). Moreover, *in vivo* metabolic flux assessment using ^13^C hyperpolarized magnetic resonance (Figure 2g) showed that *db/db* mice had 95% lower cardiac pyruvate dehydrogenase flux *in vivo*, reflecting a reduction in cardiac glucose oxidation (Figure 2h).

Remarkably, treatment with AZD1656 ameliorated cardiac metabolic dysfunction in db/db hearts. PCA analysis showed that the metabolomic profile post-drug treatment was indistinguishable between AZD1656-treated db/db hearts and controls (Figure 2a Supplementary Figure 5). The ^1^H NMR metabolomic profile was comparable between drug-treated db/db and control hearts (Figure 2c) short of persistent supra-normal elevation of succinate, an increase in NADH and a reduction in glutamine concentration. *In silico* modelling also showed improvements in cardiac metabolic flux reactions (Figure 2d), including markedly improved lipid oxidation, ROS production and hexosamine pathway utilization (Figure 2e). Our CardioNet flux analysis (Supplementary Data 1) predicts that AZD1656 treatment results in a marked reduction in anaerobic glycolysis leading to a reduction in lactate release compared to db/db hearts. One of the most notable post-drug treatment adaptations is the striking 2-fold reduction in CardioNet-calculated oxygen consumption (Figure 2f) returning it to near-control rate. Furthermore, ^13^C hyperpolarized magnetic resonance demonstrated that *in vivo* cardiac metabolic flux was improved (Figure 2h), showing increased conversion of [1-^13^C] pyruvate to ^13^C-bicarbonate, indicative of increased PDH flux.

Of note, in healthy hearts, acute AZD1656 treatment alone did not exert any metabolic or functional effects (Supplementary Figure 6). There was no difference in lipoprotein lipase (LPL) activity between control and untreated db/db hearts (control=0.84±0.13; db/db=0.66±0.09 mUnits/ml/mg), but AZD1656 treatment significantly increased LPL activity in db/db hearts (db/db=0.66±0.09 vs. AZD1656=1.34±0.13 mUnits/ml/mg, P<0.05). Furthermore, AZD1656 treatment had no impact on mitochondrial cristae density (Supplementary Figure 7).

To further study the impact of AZD1656 on cardiac lipid metabolism, we comprehensively quantified the distribution and abundance of 13 major lipid classes in heart tissues by non-targeted MS/MS lipidomic analysis (Figure 3, Supplementary Data 2). We analysed lipidomics datasets by supervised multivariate classification using partial least square-discriminant analysis (PLS-DA).

**Figure 3.**
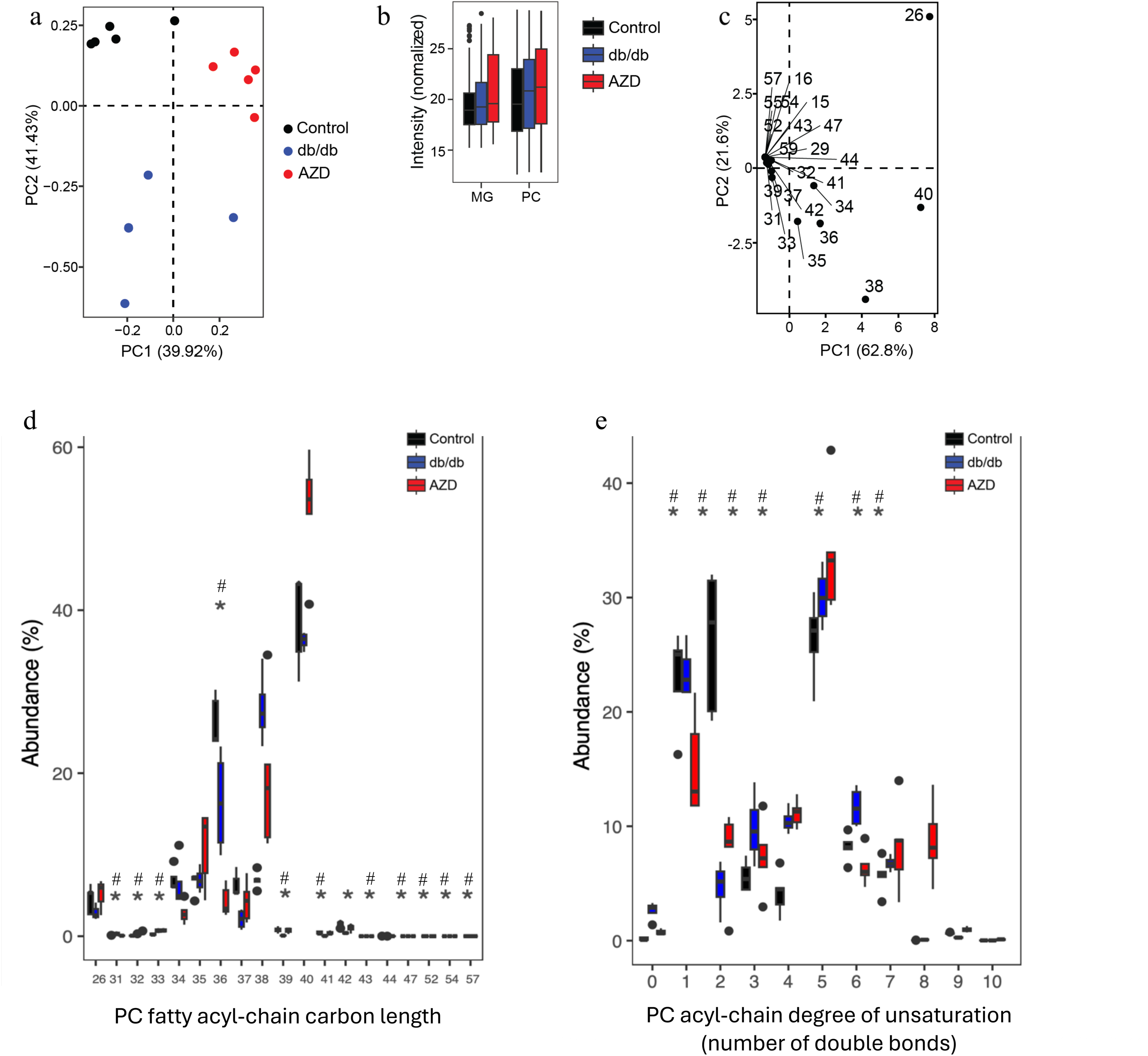
AZD1656 treatment improved cardiac lipidomic profile. **a**) Partial least square-discriminant analysis (PLS-DA) visualization of lipid abundances. Samples (n=5/group) are grouped according to biological replicates and dimensions (Dim) 1 and 2 together captured 81.35% of variability. **b**) Contribution of significantly altered phospholipid (PL) classes across biological samples. MG, Monoacylglycerols; PC, phosphatidylcholine **c**) Principal component analysis (PCA) of PC acyl chain profiles. Total carbon lengths were compared across biological samples. Dimensions 1 and 2 captured 84.4% of variability. PC(40) and PC(26) were the strongest contributors to PC alterations in treatments. **d-e**) distribution of total phosphatidylcholine (PC) fatty acyl chain length and the degree of saturation for control, db/db and AZD1656 mice. *P<0.05. control vs db/db #P<0.05 control vs AZD1656 by one-way ANOVA. P values for each comparison are provided in supplementary excel file (lipidomics comparison P values).

Among treatment-dependent separation, 81% of the total variance was captured in the first two dimensions (Figure 3a). Phosphatidylcholine (PC) and monoacyl-glycerols (MG) showed significant alterations across all three experimental groups (Figure 3b). Within the fatty acyl (FA) pools of PC, the most influential summed acyl chain lengths were identified by principal-component analysis (PCA, Figure 3c), which showed that FA26, FA38, and FA40 were dominant. Additionally, FA34, 35 and 36 contributed to PC variability. Thus, we analysed the PC fatty acyl-chain composition (carbon length) and the degree of unsaturation (number of double bonds) across all experimental groups. We found that db/db mice had decreased FA36 and FA40 abundance, while FA38 was significantly upregulated compared to control animals (Figure 3d). In contrast, treatment with AZD1656 attenuated these FA profiles towards control levels by reducing the overall contribution of FA38 and increasing the abundance of FA26 and FA40. Both db/db and AZD1656 increased the unsaturation of PC species from mono-unsaturated towards polyunsaturated fatty acyls with 4 to 5 double bonds. Untreated db/db mice had generally longer highly unsaturated PC acyl chains (6 or more double bonds) (Figure 3d). In contrast, treatment with AZD1656 caused less saturated and shorter acyl chains. The di-unsaturated to mono-unsaturated ratio was typically lower in untreated db/db hearts (0.07) compared to AZD1656 treated animals (0.2) and controls (0.129).

### AZD1656 treatment effects are not mediated by changes in systemic metabolism

In order to examine whether the AZD1656 treatment driven cardiometabolic improvements were a result of systemic metabolic alterations in db/db animals (Figure 4a), extensive analyses of blood, liver, adipose tissue and skeletal muscle were performed. AZD1656 treatment did not improve obesity (body weight, Figure 4a, b), hyperglycaemia (Figure 4c, Supplementary Figure 8), hyperinsulinemia (Figure 4d) or insulin resistance (adiponectin, Supplementary Figure 8). Whilst AZD1656 treatment significantly reduced the circulating free fatty acid concentration (Figure 4e), the remnant circulating metabolite profile was not improved (Figure 4 f,g, Supplementary Figure 8). AZD1656 treatment did not improve the diabetic liver phenotype (Figure 4h, alanine transferase, alkaline phosphatase, Supplementary Figure 8) or metabolism (Liver ^1^H NMRs metabolomics Figure 4i,j, Supplementary Figure 9a). AZD1656 treatment led to a minute increase in lean muscle mass vs db/db (Figure 4k) but had limited effect on the skeletal muscle metabolome (Figure 4 l, m, Supplementary Figure 9b). Furthermore, treatment with AZD1656 did not reduce whole body fat mass (Supplementary Figure 10a) nor alter the lipid composition of the adipose tissue (Supplementary Figure 10).

**Figure 4.**
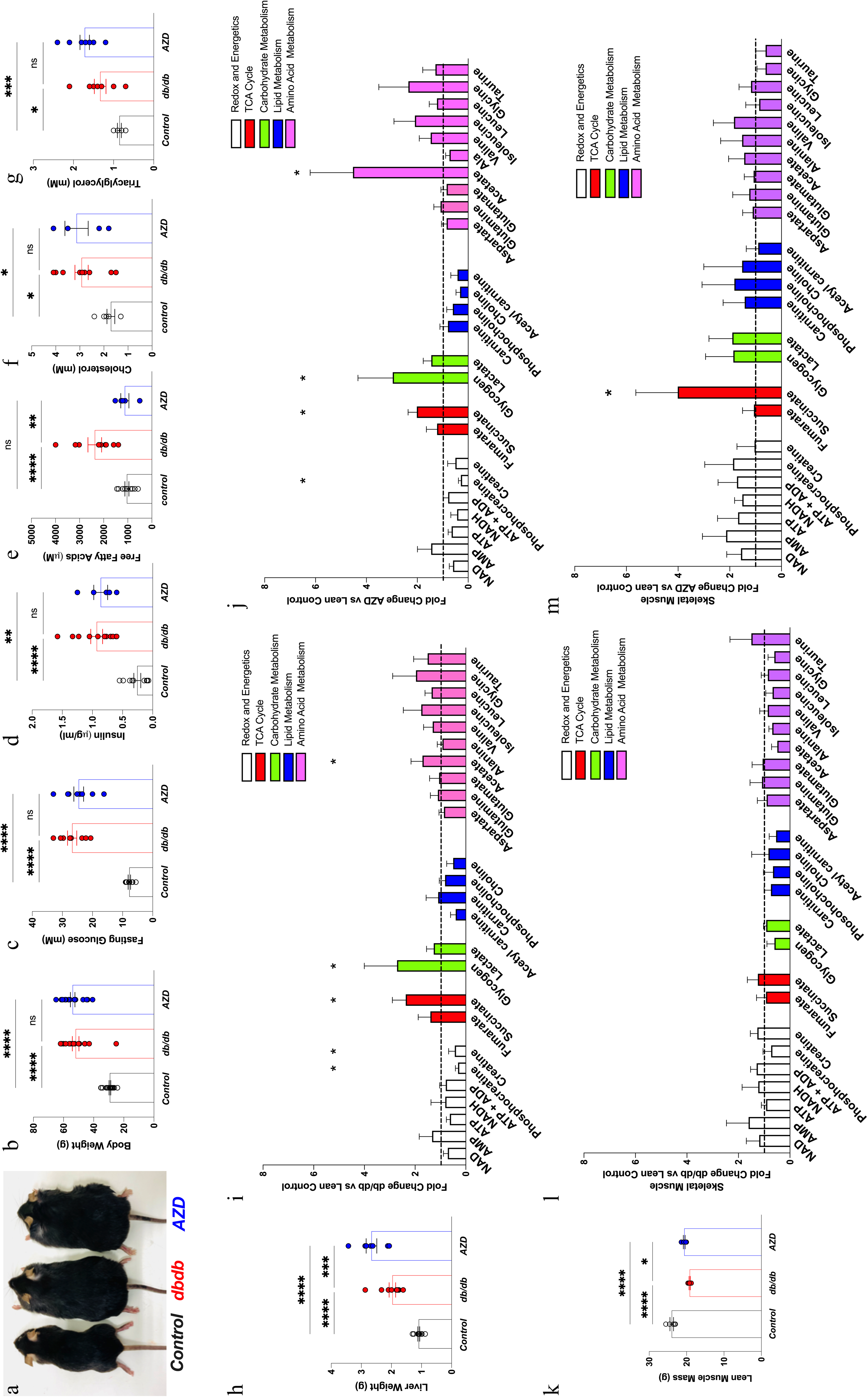
The systemic effect of AZD1656 treatment in db/db. **a)** Representative images of control, db/db and AZD treated db/db mice from the study **b**) Body weight (Control n=23, db/db n=17, AZD n=20) **c**) Fasting plasma glucose (n=9/group) **d**) plasma insulin (Control n=10, db/db n=11, AZD n=6) **e-g**) circulating lipid metabolism constituents [free fatty acids (Control n=11, db/db n=9, AZD n=5), cholesterol (Control n=7, db/db n=10, AZD n=5), triacylglycerol (Control n=7, db/db n=11, AZD n=9). Liver phenotype profiling **h**) Liver weight (Control n=12, db/db n=11, AZD n=7) **i**) Fold change db/db vs control ^1^H NMR metabolomic spectroscopy profiling **j**) Fold change AZD vs control ^1^H NMR metabolomic spectroscopy profiling (Control n=6, db/db n=7, AZD n=7) Skeletal Muscle phenotype (Control n=6, db/db n=6, AZD n=5) **k**) lean muscle mass by body composition analysis (n=5/group) **l**) Fold change db/db vs control ^1^H NMR metabolomic spectroscopy profiling, **m**) Fold change AZD vs control ^1^H NMR metabolomic spectroscopy profiling, (n=5/group) *P<0.05, **P<0.01, ***P<0.001 multiple group comparison by ANOVA, two-tail comparison by student t-test. *Note:* Parameters measured over the course of 6-year study in multiple cohorts thus sample size varied.

### AZD1656 treatment reduces T-cell mediated cardiac inflammation and fibrosis in dbCM

Untreated db/db mice were characterized as having low-grade systemic inflammation, observed from alterations to the circulating cytokine profile (Figure 5a, Supplementary Figure 11). There was an increase in IGF BP-1, LDL-R, PD-ECGF / thymidine phosphorylase, leptin and osteoprotegerin.

**Figure 5.**
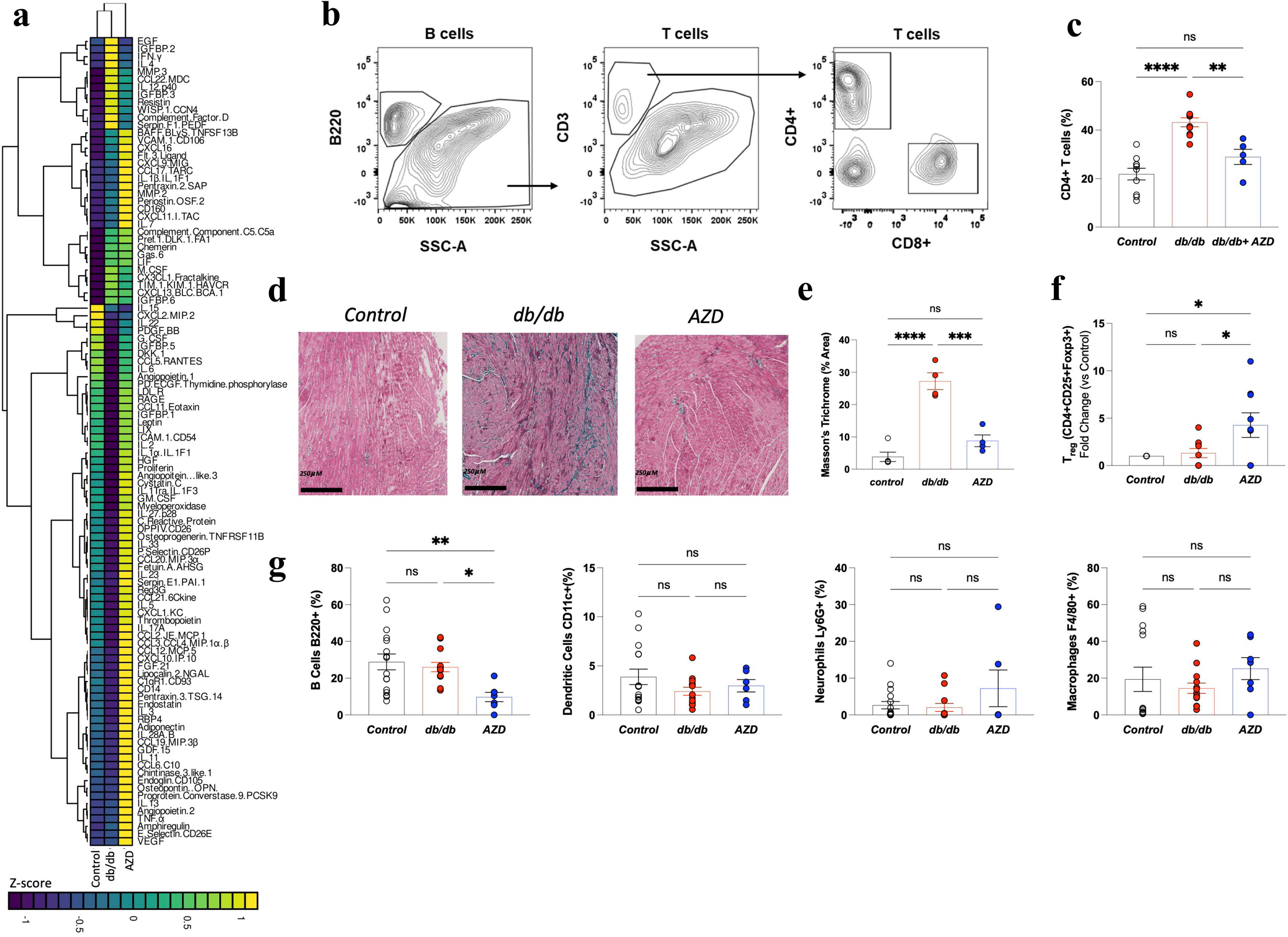
AZD1656 treatment reduces T-cell mediated cardiac inflammation and fibrosis in dbCM. **a)** Circulating XXL cytokine plasma panel heatmap summary (Control n=10, db/db n=12, AZD n=10 **b)** Representative FACS contour plots **c**) Relative frequency of myocardial CD4+ T cells quantified by FACS **d**) Masson trichrome stained representative cardiac cross sections **e**) quantification of Masson trichrome fibrosis staining intensity (Control n=5, db/db n=4, AZD n=4) **f**) Relative frequency of myocardial Treg cells (CD4+CD25+FoxP3) (n=9/group); **g)** dendritic cells, neutrophils and macrophages (controls n=8, db/db n=8, AZD n=5) *P<0.05, **P<0.01, ***P<0.001.Multiple group comparison by ANOVA.

A significant decrease was observed in circulating anti-inflammatory mediators IL-10, IL-4, growth repair factors EGF, PD-ECGF, WISP-1, Serpin F, matrix remodelling enzymes MMP-3, WISP-1, chemokines CX3CL1, CXCL16, CD40, immune cell regulators M-CSF, complement factor D, IL-12p40, adipokines/metabolic signal resistin (Figure 5a, Supplementary Figure 11).

Unlike in resolving acute cardiac inflammation in which infiltrating Tregs are increased^47^, cardiometabolic and functional derangements in db/db were concomitant with unregulated T cell-mediated cardiac autoimmunity (Figure 5b,c) and fibrosis (Figure 5d,e) without Treg increase (CD4+CD25+FoxP3+, Figure 5f). In terms of infiltration of other inflammatory cell types, there was no evidence of increased presence of B cells (B220), Ly6C-lo monocytes, CD11c+ dentritic cells, neutrophils Ly6G+ or macrophages F4/80 (Figure 5g). In pre-diabetic, high-fat–fed C57BL/6 mice displaying obesity, insulin resistance and early cardiac metabolic derangement preceding overt T2D, no myocardial infiltration by T cells or other inflammatory cells (macrophages, dendritic cells, neutrophils, B Cells) was observed (Supplementary Table 2, Supplementary Figure 12).

AZD1656 markedly attenuated cardiac inflammation in db/db mice. Treated hearts exhibited reduced interstitial fibrosis (Fig. 5d,e) and diminished T-cell infiltration (Fig. 5c), alongside a pronounced increase in Foxp3□ regulatory T-cell recruitment (Figure 5f), evidence of enhanced Treg mobilization and a more favourable myocardial immune milieu^48^. AZD1656 treatment did not induce global CD4□ T-cell expansion, as evidenced by unchanged splenic CD3□ T-cell abundance, consistent with the absence of systemic lymphoproliferative effects (Supplementary Figure 13). While AZD1656 did not alter dendritic cell, neutrophil or macrophage trafficking (Fig. 5g), it significantly decreased B-cell accumulation within the myocardium (Fig. 5g).

In terms of soluble cytokine profile, there was a marked change post-AZD1656 treatment (Figure 5a, Supplementary Figure 11). Pharmacological intervention normalised plasma concentrations of IL4, BAFF, CCL6, FGF-21, GDF-15, coagulation factor III, CXCL16, FGF acidic, Gas6, IL-12p40, M-CSF, CXC3CL1, MMP3, decreased concentrations of IL10, serpin F1, resistin, CD40, EGF and complement factor D and increased concentrations of CCL20, CCL22,CXCL1/KC, MMP-9, Serpin E1, P-selectin, myeloperoxidase, PDGF-BB, IFGBP-1, LDL-R, PD-ECGF, leptin and osteoprogenerin (Figure 5a, Supplementary Figure 11).

### Treatment with AZD1656 normalizes the expression of genes regulating key intracellular pathways

Development of dbCM in db/db mice was characterized by significant alterations in cardiac gene expression [1379 differentially expressed genes (DEGs) db/db vs control, 503 genes db/db vs AZD1656, 1480 genes Control vs AZD1656, Figure 6a]. G profiler gene enrichment analysis (Supplementary Figure 14) shows dbCM caused extensive alterations in the expression of genes governing key aspects of cardiac function: molecular functions, biological processes, cellular components, protein complexes as well as biological pathways. Analysis of the 30 most upregulated genes in db/db vs control hearts shows that 19 out of the top 30 over-expressed genes are pro-inflammatory mediators (Figure 6b).

**Figure 6.**
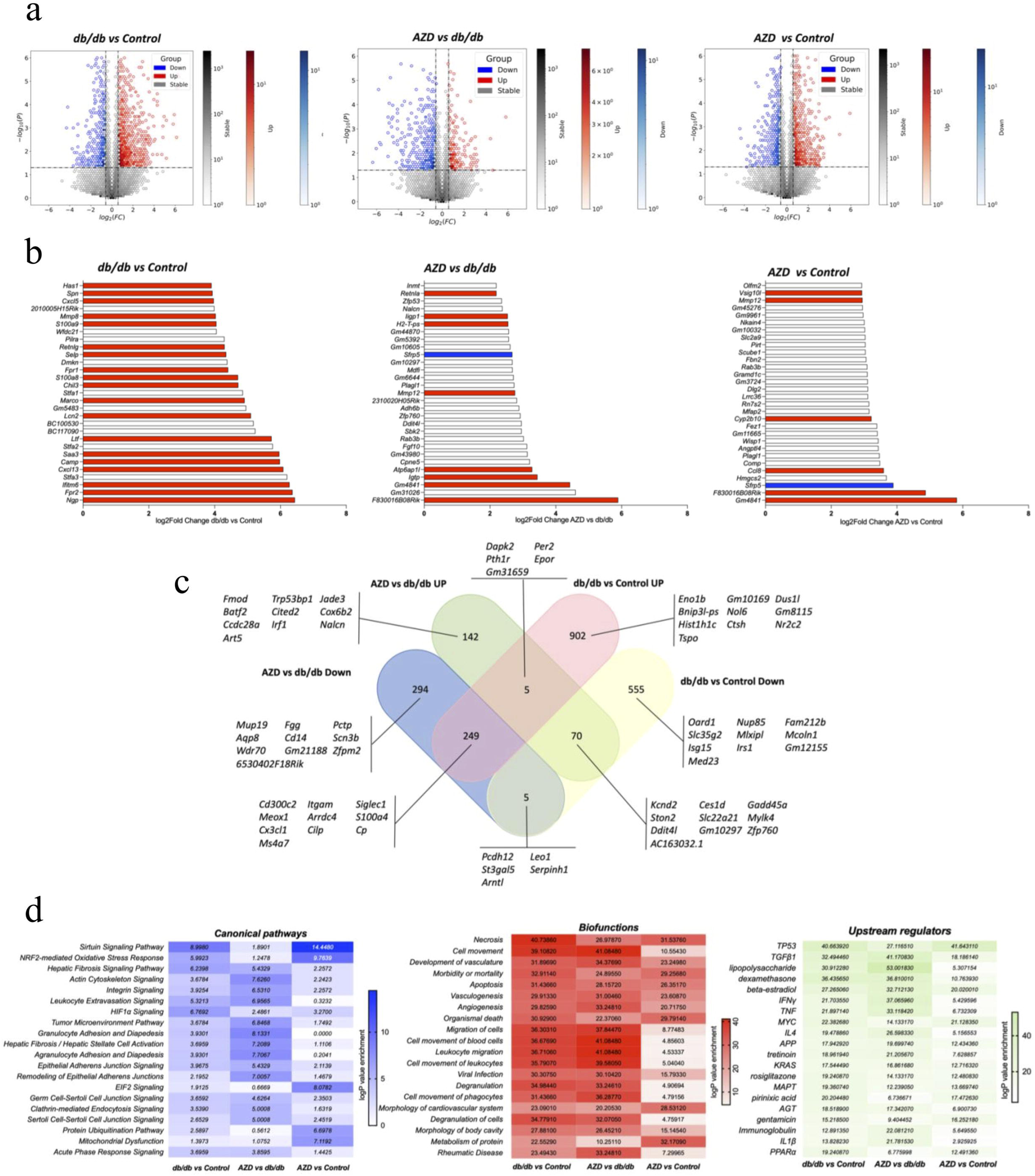
Treatment with AZD1656 improves the expression of genes regulating key intracellular pathways. Visualisation of differentially expressed genes (DEGs) using hexbin plot summaries. X axis values are log2fold Change vs y axis - log 10 (P value) of **a**) db/db vs Control, AZD vs db/db, AZD vs Control. Y axis vertical cut off log2 (1.5) ± 0.585 and x axis horizontal cut off -np.log10(0.05) ∼ 1.301 **b**) Top 30 most upregulated DEG in db/db vs Control, AZD vs db/db, AZD vs Control. Red bars pro-inflammatory regulators, blue bar-anti-inflammatory regulators. **c**) The Venn diagram of overlapping homologous DEGs among the three data sets (Control, db/db and AZD). Threshold was set to be FC > 1.5 (or <1/1.5) and P < 0.05. Sample genes ( increased and decreased) from each of the subsets are given: abs_diff_log2FC = abs(log2FoldChange_AZ_vs_dbdb - log2FoldChange_dbdb_vs_CTRL) **d**) Heatmap plots of - log p value of DEGs resulting from hypergeometric test and Benjamini-Hochberg p-value correction to identify all ontology and pathway terms that contain a greater number of genes in common with an input list than expected by chance using Qiagen IPA software^38^. This is expressed as a - log p value in the heatmaps. All RNA seq data presented in this figure is based on n=6 hearts/group.

*Nlrp3* gene inflammasome levels were markedly elevated, showing a 2.5-fold increase in db/db compared with controls (RNA sequencing data, Array Express accession E-MTAB-13849). This upregulation suggests enhanced assembly of the inflammasome complex, which likely drives increased caspase-1 activation and subsequent IL-1β and IL-18 release, promoting a pro-inflammatory myocardial environment.

Further Venn diagram analysis (Figure 6c) identified overlapping homologous DEGs of db/db, control and AZD1656-treated db/db hearts. The Venn diagram analysis of gene homology between the groups highlights the upregulation of the inflammation mediating genes (*Frp2, Ifitm6, Camp, Ltf, Ngp, Saa3*) including leukocyte migration mediator *Selp* in db/db versus lean controls (Figure 6c). Pathway analysis further identified genetic hallmarks of dbCM and has shown the upregulation of canonical pathways, biofunction regulators and upstream regulators (Fig 6d) of inflammation, apoptosis, T-cell migration/infiltration, inflammatory signalling and metabolism (fats and carbohydrates) (Supplementary Figures 14-17).

AZD1656 treatment of T2D markedly changed the altered gene expression profile of T2D hearts (Figure 6 a,b) as treatment reversed pathologically high gene expression in db/db hearts compared to untreated (503 genes, AZD1656 vs db/db Figure 6b). Out of the 30 most expressed DEGs in AZD1656 vs db/db hearts and AZD1656 vs Control hearts only 8 (AZD1656 vs db/db) and 6 (AZD1656 vs Control) are pro-inflammatory. Of note, AZD1656 treatment also increased expression of *Sfrp5* (frizzled-related sequence protein 5), a mediator of protection against inflammation and apoptosis via inhibition of Wnt5a/JNK pathway^49^. AZD1656 treatment reduced the number of shared, upregulated inflammation mediators in AZD1656 versus Control to only 3 genes (Venn diagram, *GM4841, F830016B8Rik, Sfrp5*, Figure 6c). Pathway analysis reveals that drug treatment resulted in the opposite pattern of myocardial pathway gene expression: for instance, the allograft rejection pathway that showed the biggest increase in log fold change in expression in db/db vs control is the least expressed in db/db vs AZD1656 (Supplementary Figure 14). These gene expression changes driven by AZD1656 treatment also included a reduction in the expression of genes in reactive oxygen species production pathways, metabolism of lipids and multiple inflammation mechanisms and signals (Figure 6d).

In addition, treatment with AZD1656 targeted and improved a whole series of pathways involved in biological functions (Figure 6d) including necrosis, cell movement, allograft rejection, apoptosis, leukocyte migration, signalling pathways (sirtuin, Nrf2, hepatic signalling, leukocyte migration and signalling, Figure 6d, Supplementary Figure 15). Furthermore, AZD1656 treatment impacted a range of upstream regulators including improvement of pro-inflammatory signals (TGFβ1, IFNγ, IL4, PPARα Supplementary Figure 14).

Crucially, the mechanistic impact of the AZD1656 treatment on db/db hearts was shown by complete amelioration of the elevated *Nlrp3 expression* (RNA sequencing data, Array Express accession E-MTAB-13849) as well as dysregulated leukocyte extravasation signaling governing T-cell infiltration of the myocardium (change of signaling pathway intermediate components from red to green, detailed Kegg pathway in Supplementary Figure 15) as well as of two key metabolic regulator pathways: HIF1α (Supplementary Figure 16) and NRF2-mediated Oxidative Stress Response (Supplementary Figure 17).

## Discussion

Given the previously proposed immunomodulatory effect of AZD1656^5,6^, we investigated the possibility that this treatment would improve cardiac inflammation and in turn attenuate cardiac remodelling in dbCM. Systemic T-cell mediated inflammation has been extensively phenotyped in T2D^50^ and other chronic metabolic stress states^1,51^, however cardiac inflammation in T2D remained to be identified and profiled.

As previously observed, 20-week old db/db mice were defined by the phenotypic features representative of human dbCM: obesity, hyperglycaemia and impaired diastolic function (*in vivo* and *ex vivo*) (summarised in Figure 7). Analysis of 20-week db/db diabetic hearts showed altered T-cell dynamics accompanied by an increase in systemically altered pro-inflammatory cytokine production and fibrosis, which was concomitant with altered cardiac metabolism: demonstrated through *in vivo* substrate utilisation (hyperpolarized ^13^C magnetic resonance spectroscopy), energetics, oxidative phosphorylation, and amino acid metabolism (^1^H NMR spectroscopy).

**Figure 7.**
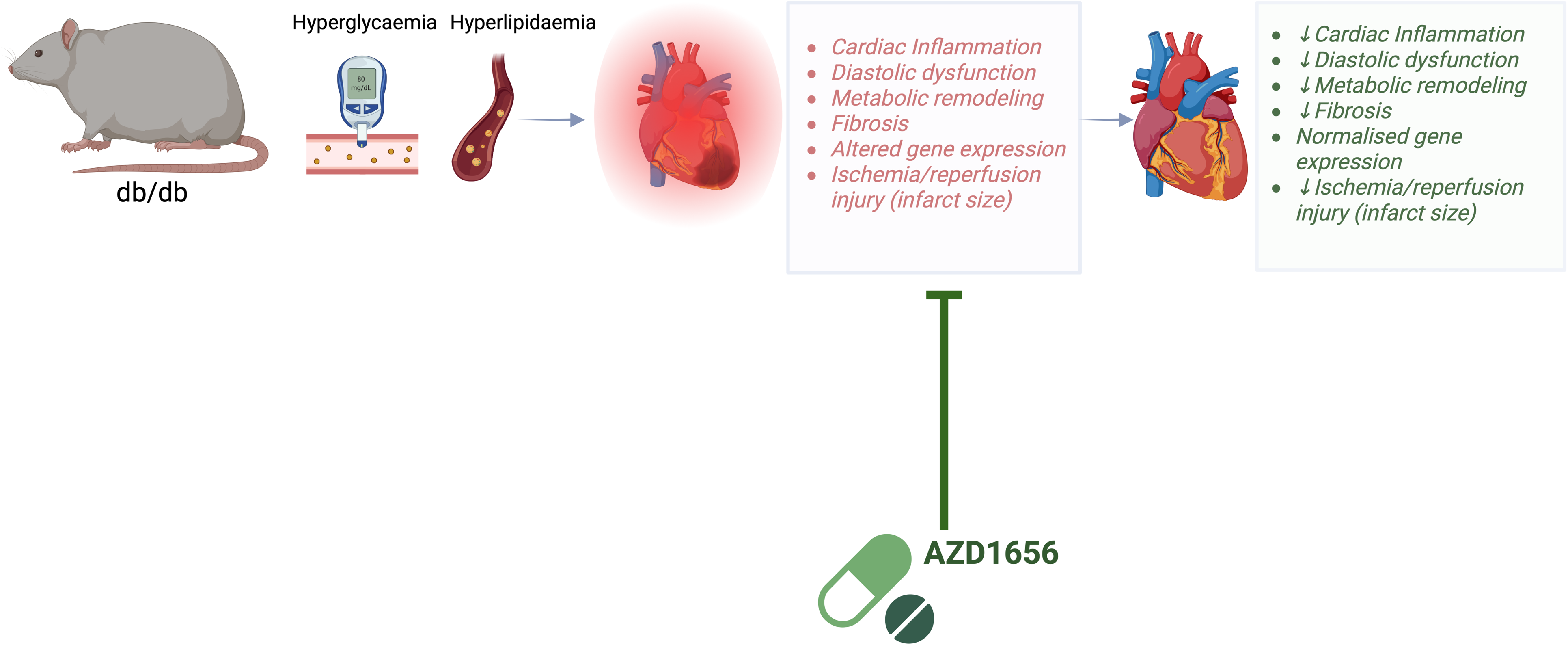
Summary of the key study findings. AZD1656 treatment may represent a new therapeutic concept for the treatment of dbCM-the use of small molecules targeting immunometabolism to reduce cardiac inflammation in the diabetic heart. Figure generated with Biornder.com

Remarkably, 6-weeks of pharmacological treatment with AZD1656 attenuated the functional and metabolic remodelling observed in the untreated diabetic heart, resulting in improved post-ischemic functional recovery and reduction in ischemic damage.

One of the key challenges in dbCM is the loss of metabolic flexibility^52^. However, AZD1656 drug treatment achieved significant improvements in the cardiac metabolomic profile and flux as observed from both *in vivo* and *in silico* metabolic assessment. These changes are suggestive of improved substrate utilization flexibility. Furthermore, AZD1656 pharmacological intervention in db/db mice resulted in a 68% reduction in cardiac oxygen consumption. Increased myocardial oxygen consumption and O_2_ wasting due to uncoupled mitochondrial function have been shown to reduce cardiac efficiency in T2D ^53,54^. Thus, the reduction in oxygen consumption post-AZD1656 treatment is critical for the improvement in cardiac function and for the reduction in ischemia-reperfusion damage.

dbCM in db/db mice was also characterized by differences in the fatty acyl chain composition of glycerolipids such as phosphatidylcholine as well as monoacyl-glycerols. AZD1656 treatment significantly improved the cardiac phospholipid composition suggestive of improved lipid metabolism as well as cellular membrane and mitochondrial integrity^55^ ^56^. Our studies demonstrated that pharmacologic interventions that attenuate obesity-induced alterations in phospholipid composition and homeostasis positively impact cardiac metabolism and function. Bulk cardiac tissue RNA sequencing analysis showed that AZD1656 treatment significantly improved the gene expression profile including genes governing key metabolic and inflammatory response pathways (ie. NLRP3, HIF1α, PGC1α, apoptosis, necrosis, leukocyte extravasation, fatty acid metabolism, oxidative phosphorylation).

Experimental AZD1656 treatment outcomes could have been achieved via two potential mechanisms of action: by improved glycaemic control (original drug indication) and/or by modulation of the immune response^6^. Since no differences in glycaemic control were detected between groups, it is unlikely that the differences in outcomes were a result of the glucose lowering effect of AZD1656.

T-cell inflammation has been previously linked with the development of heart failure in both animal disease models and humans^57^. T-cell costimulatory blockade, using the rheumatoid arthritis drug abatacept, significantly reduced the severity of cardiac dysfunction in HF. This therapeutic effect occurred via the inhibition of activation and cardiac infiltration of T-cells and macrophages, leading to reduced cardiomyocyte death.^57^ The immunophenotyping data here suggested that AZD1656 treatment induced an immunomodulatory effect in db/db mice via improved Treg infiltration, reduction in pro-inflammatory, CD4+T-cells and the resultant reduction in fibrosis. This is consistent with rebalancing of the immune response observed in the ARCADIA trial in COVID19 T2D patients^6^. Treg migration is metabolically demanding and fuelled by glucokinase-regulated glycolysis^5^. This has been shown in human carriers where a loss of function gene polymorphism in GCK regulatory protein (GKRP) causes Tregs, rather than conventional T-cells, to display altered motility *ex vivo*^5^. Whilst glycolysis fuels the migration of other T-cell subtypes including pro-inflammatory T effector cells, their migration is hexokinase I (HKI) rather than glucokinase (HKIV) dependent^58^. Thus, GCK activator AZD1656 selectively enhanced Treg migration and resultant myocardial infiltration.

Increased infiltration of Tregs into the db/db myocardium exerts multi-faceted cardioprotective effects by simultaneously modulating the immune landscape, metabolic efficiency, and tissue remodelling. Tregs secrete key anti-inflammatory cytokines including IL-10 and TGF-β, which suppress pro-inflammatory macrophage activation and effector T cell infiltration, thereby attenuating the chronic low-grade inflammation characteristic of pathologies such as HF^59,60^. Treg-secreted anti-inflammatory cytokines limit NLRP3 inflammasome assembly^61^ which would account for normalised *Nlrp3* gene expression in AZD16556-treated db/db hearts.

The Treg-mediated immunosuppressive milieu directly translates into a reduction in maladaptive fibrotic remodelling through inhibition of cardiac fibroblast activation and downregulation of TGF-β/Smad signalling pathways^62^. Furthermore, this immunomodulation has the potential to restore insulin signalling pathways, thereby improving cellular glucose uptake and mitochondrial function^63^. In organs with high metabolic demand such as the heart, enhanced Treg presence also promotes expression of key metabolic regulators such as PPARγ further improving overall metabolic homeostasis.^64^ Beyond immunoregulation, Tregs influence vascular integrity by promoting endothelial quiescence and angiogenic support, improving myocardial perfusion in the context of microvascular dysfunction^59^. In models of cardiac ischemia/reperfusion injury, Treg-derived amphiregulin and IL-33 signalling not only promote tissue regeneration but also limit infarct expansion^65^. This could account for the observed reduction in infarct size in AZD1656-treated db/db hearts. Tregs further support cardiomyocyte survival by enhancing IL-10-mediated activation of STAT3 signaling^66^. Collectively, these findings position Tregs as potent mediators of immunometabolic resilience in dbCM.

In addition to the improved T-cell inflammatory profile, AZD1656 also caused a significant decrease in B cell infiltration versus untreated db/db. Increased B cell infiltration has been show to increased antibody (IgG3) production that aids the recruitment and activation of a variety of innate and structural cell populations: neutrophils, fibroblasts, macrophages and T-cells^67^. As AZD1656 boosts Treg infiltration and suppresses CD4+ T-cells, the myocardial environment shifts from pro-inflammatory to immunoregulatory. Tregs secrete IL-10 and TGF-β, which downregulate key chemokines and endothelial adhesion molecules (e.g., CXCL13, VCAM-1) that ordinarily recruit B-cells to inflamed tissue.

At the same time, the reduction in CD4+ deprives B-cells of essential survival and activation signals such as IL-21 and CD40L, impairing their retention and viability in cardiac tissue. Finally, by suppressing NLRP3 inflammasome activation and its downstream cytokines IL-1β and IL-18, AZD would further diminish the pro-B-cell milieu. Consistent with findings that BAFF levels rise after B-cell depletion, and given the reduction of cardiac B cells we observe with AZD1656, it is reasonable to expect a corresponding decrease in peripheral B-cell numbers following treatment^68^.Together, these mechanisms converge to lower myocardial B-cell numbers while restoring a balanced, protective immune state in db/db hearts. Given that B cells have been linked to the development of inflammation and maladaptive cardiac remodelling in HF,^67^ immunomodulatory therapeutic strategies that target B cells and their products could be an effective novel treatment tool. Our study focused on heart-resident CD4+ T cells, so we did not assess peripheral blood or lymphoid tissues. While sequencing the systemic TCR repertoire could provide complementary insights, it was beyond the scope of this cardiac-centric investigation.

AZD1656 delivers a finely tuned suppression of diabetogenic inflammation and remodelling via plasma cytokine rebalancing. Pharmacological intervention led to a coordinated increase in a suite of cytokines implicated in immune regulation, vascular remodelling, and metabolic adaptation, collectively suggesting reversal of immune suppression and reactivation of cardiac repair pathways. Key chemokines such as CCL20, CCL22, and CXCL1/KC promote the recruitment of Tregs fostering a reparative immune environment and reducing chronic inflammation^69,70^. CCL20 and CCL22 are ligands for CCR6 and CCR4, respectively, both expressed on Tregs and implicated in their trafficking to inflamed tissue. Upregulation of BAFF and IL-12p40 supports tolerogenic dendritic cell maturation and T cell differentiation, consistent with immune modulation rather than proinflammatory activation^71^. Alteration in circulating IL-12 is indicative of less T-cell effector function and DC activation^33^. Increased FGF-21 and GDF-15 are markers of systemic metabolic rescue and mitochondrial stress tolerance, both shown to be cardioprotective under diabetic conditions^72,73^. Elevated MMP-9 and Serpin E1 reflect dynamic matrix turnover and remodelling, while enhanced PD-ECGF, FGF acidic, and Gas6 suggest restoration of angiogenic signalling and endothelial repair^74,75^.

The AZD1656 triggered induction of P-selectin, CXCL16, and myeloperoxidase likely represents controlled inflammatory engagement, contributing to immune surveillance and clearance. Notably, metabolic and vascular stress mediators such as leptin, resistin, LDL-R, and osteoprotegerin (OPG) were also elevated, which may reflect attempts at metabolic compensation and vascular remodelling. Increased IGF BP-1 may indicate improved IGF signalling balance or a response to restored metabolic demand^76^.

Systemic metabolomic profiles of blood, liver, adipose tissue, skeletal muscle were largely unaffected thus any beneficial effects observed in db/db treated hearts were not driven by marked improvements in systemic peripheral tissue metabolism (liver, muscle). We have, however, observed a significant reduction in the circulating fatty acid concentration with an accompanying increase in body fat content post-AZD1656 treatment. This could have contributed to the improvement of myocardial metabolism and inflammation. Reduced circulating fatty acids would have affected myocardial metabolism, specifically PDH flux *via* fatty acid regulation of PDK4^77,78^ and reduced systemic pro-inflammatory T-cell proliferation^79^

The observed reduction in circulating free fatty acids may reflect the combined effects of AZD1656 on β-cell and hepatic glucokinase activity, leading to systemic reorganization of lipid metabolism^80–82^. In models with preserved β-cell function, glucokinase activation enhances glucose-stimulated insulin secretion and secondarily increases LPL activity through hyperinsulinemia^83^. However, db/db mice are already profoundly hyperinsulinemic with limited β-cell capacity; therefore, the increase in cardiac LPL activity and altered lipid milieu following AZD1656 treatment most likely reflect hepatic substrate-driven signaling rather than additional insulin release.

Consistent with this interpretation, AZD1656 did not exacerbate hypertriglyceridemia-plasma triglycerides remained elevated but unchanged-while circulating free fatty acids declined significantly, indicative of predominantly hepatic, insulin-independent effects. Hepatic glucokinase activation may enhance glycolytic flux and lipogenesis, maintaining VLDL-triglyceride output without further elevating plasma triglycerides. The resulting lipid redistribution could stabilize myocardial LPL^84^. However, despite increased cardiac LPL activity, fatty-acid utilization was not augmented; rather, AZD1656 normalized PDH flux and restored a more balanced cardiac metabolic profile, suggesting improved glucose oxidation and metabolic flexibility via hepatic-cardiac metabolic cross-talk. Given negligible cardiac GCK expression and no direct metabolic effect in perfused hearts, enhanced myocardial LPL activity likely represents a secondary adaptive response, complementing the drug’s immunomodulatory actions. Supporting this, individuals carrying the GCKR P446L loss-of-function variant-which increases hepatic glucokinase activity-exhibit similar alterations in systemic lipid metabolism and enhanced Treg motility^5^.

If AZD1656 benefits truly depend on modulating Tregs, why do we observe its anti□inflammatory effects exclusively in the heart? Ultimately, why T cells preferentially home to the diabetic myocardium despite systemic dysregulated metabolism remains an open, almost philosophical, question that transcends current mechanistic understanding. One of the key characteristics of dbCM is an increase in fibroblasts which have been shown to act as antigen presenting cells^85^ driving myocardial T-cell infiltration and inflammation^86^. In db/db mice, potentially there are four converging factors render the myocardium uniquely immunogenic: (1) chronic hyperglycemia and lipotoxicity selectively up-regulate CXCL10, CCL2, VCAM-1 and ICAM-1 in cardiomyocytes and coronary endothelium, establishing “docking sites” for lymphocytes^87^; (2) metabolic stress generates AGE- and oxidized-lipid neoantigens that are robustly presented on myocardial MHC-I/II, locally activating CD4□ T-cell responses; (3) heightened TLR2/4 signaling in resident macrophages and cardiomyocytes fuels IL-1β, IL-6 and TNFα release, amplifying antigen presentation and chemotaxis^88^; and (4) by contrast, skeletal muscle, liver and adipose tissue maintain lower basal levels of cardiac self-antigens and inducible adhesion molecules, failing to create the chemokine gradients or MHC up-regulation needed for substantial T-cell homing. Together, these factors create a “perfect storm” in diabetic hearts: enhanced antigen presentation, chemokine expression and endothelial adhesion molecule display converge to selectively recruit and retain T-cells in the myocardium, driving inflammation and dbCM phenotype. Together, these cardiac-restricted antigenic and adhesion cues explain the selective myocardial infiltration and thus the heart-specific anti-inflammatory effects of AZD1656-targeted Tregs in diabetic cardiomyopathy. Future studies using CD4□ T cell-deficient mice or antibody-mediated depletion would be valuable in delineating the mechanistic contribution of CD4□ T cells to dbCM. These studies are currently under consideration for follow-up work.

Despite these promising results presented in this work, questions remain relating to whether the changes in cardiac metabolism or immune response are the therapeutically exploitable culprits in triggering the dbCM pathology. Specifically, what is the temporal sequence of events: is mitochondrial dysfunction an early driver, or does inflammation occur first?

To help to understand the sequence of events driving diabetic cardiomyopathy, we established a spectrum of cardiac dysfunction by inducing obesity and insulin resistance in mice via a high□fat diet, using age□, sex□, and strain□matched controls to db/db^13^. In this pre□diabetic model, we observed clear cardiac metabolic derangements before any evidence of myocardial immune cell infiltration. These findings indicate that systemic metabolic disturbances (hyperglycemia, glucose intolerance, obesity) and intrinsic mitochondrial dysfunction are primary events that precede, and likely trigger, subsequent cardiac inflammation in established T2D such as that seen in db/db mice. This would be in agreement with a recently proposed model where mitochondrial dysfunction itself is a potent pro□inflammatory trigger^89^. When oxidative phosphorylation falters, excess mitochondrial ROS and oxidized mtDNA leak into the cytosol, where they directly engage and activate the NLRP3 inflammasome. This drives caspase□1-dependent maturation of IL-1β and IL-18, which in turn amplify NF-κB signaling and downstream production of TNF and IL-6. Simultaneously, released mitochondrial lipids such as cardiolipin act as danger□associated molecular patterns, further escalating cytokine release^89^. Thus, broken mitochondria are not silent bystanders but the very “damage sensors” that convert metabolic stress into chronic, low-grade inflammation^89^.

A growing body of evidence demonstrates that pharmacological augmentation of Tregs confers potent anti□inflammatory and plaque□stabilizing effects in models of cardiovascular disease. Agents as diverse as rapamycin and mycophenolate mofetil expand CD4□FoxP3□ Tregs while concurrently depleting pro□inflammatory effector T cells and antigen□presenting cells, thereby reducing vascular inflammation and matrix metalloproteinase activity^90^. Statins, aspirin, and PPAR□γ agonists (pioglitazone) similarly restore the Th17/Treg balance, elevate IL-10 and TGF-β, and promote endothelial homeostasis. Even microbiome□derived metabolites (indole propionic acid) and micronutrients (vitamin D□, amygdalin) have been shown to increase Treg frequencies in hypertensive and atherosclerotic models^90^. Collectively, these interventions validate Treg modulation as a unifying strategy to attenuate chronic vascular inflammation and improve metabolic resilience in cardiometabolic disease. However, the ARCADIA trial outcome has introduced a novel immunometabolic therapeutic concept: pharmacological targeting of the endogenous immune cells to turn them into the therapeutic agents within the body^6^. Previously, this drug development approach, whether it was used in exogenously treated or engineered cells has failed^6,91–93^. Our study results contribute to the growing field of cardio-immunology as they suggest that AZD16565 could be used as a novel immunomodulatory drug to attenuate cardiometabolic dysfunction in type 2 diabetes as well as the whole host of sterile-inflammation pathologies.

## Data Availability

RNA sequencing data is available (open access) on Array Express accession E-MTAB-13849.

All datasets supporting the findings of this study, including ex vivo function, morphology, biochemical assays, ^1^H NMR metabolomic analysis, echocardiographic, flow cytometric, and transcriptomic data are available on the Dryad Digital Repository: http://datadryad.org/share/LINK_NOT_FOR_PUBLICATION/RHY3R9GZFltMKv1thb_5dEustRJ9b-qQH4-8GjY0sIU

All scans of infarcted hearts corresponding to Langendorff perfusion and TTC staining analyses are provided in Supplementary File.

Detailed results from the Cardionet in silico analyses and all lipidomic mass spectrometry data are included in the Supplementary Data Files.

Furthermore, the data that support the findings of this study are also available from the corresponding author (Prof D Aksentijevic) upon request.

## Manuscript Correspondence

Any correspondence regarding me manuscript needs to be directed to Prof Dunja Aksentijevic

## Acknowledgements

DA acknowledges Mrs Irineja Cubela (Barts Cancer Institute, QMUL), Dr Harold Toms, Dr Roberto Buccafusca, Dr Nasima Kanwal (School of Physical and Chemical Sciences, QMUL), Dr T. Eykyn (King’s College London), Dr Giulia Mastroianni (School of Biological and Chemical Sciences), Mr Ahmad Hjiej Andaloussi for technical assistance. DA is the recipient of the Wellcome Trust Career Re-Entry Fellowship (221604/Z/20/Z) and acknowledges the following funding sources: British Heart Foundation Accelerator Award fellowship (AA/18/5/34222), Diabetes UK Grant (19/0005973), Barts Charity Grant (MRC 0215, G-002145), Astra Zeneca UK (study agreement ref 3853), School of Biological and Behavioural Sciences QMUL (HEFCE lectureship). SMH was funded by Diabetes UK 19/0006057, CGK by Barts and the London Charity MGU0536. DJT was funded by British Heart Foundation Senior Basic Science Research Fellowships (FS/14/17/30634 and FS/19/18/34252). SA was funded by the Pelly-Carys Bannister Scholarship, Somerville College, University of Oxford. AK was supported by the National Institute of Health (NIH) (R00-HL-141702 to A.K.), the Leukemia Research Foundation (New Investigator Award, Grant No. 941997) and the Cedars-Sinai Cancer Center through the 2022 Cancer Biology Program CardioOncology Award. MRB acknowledges funding from the NIHR BRC at Barts and The London School of Medicine and Dentistry. LJA acknowledges British Heart Foundation Programme Grant RG/12/4/29426. CO’R was sponsored by Barts and The London School of Medicine and Dentistry, Queen Mary University of London. JC was supported by a Career Development Fellowship of Versus Arthritis UK (22855). JJM acknowledges support of a faculty grant from the Novo Foundation (NNF21OC0068683). This work was supported by the Medical Research Council UK (MC_UU_00028/4) and by a Wellcome Trust Investigator award (220257/Z/20/Z) to MPM. This research was supported by the MRC Mouse Biochem Lab, Cambridge [Medical Research Council (MC_UU_00014/5)]. Figure 1a and Figure 7 generated with Biorender.com. Sofia and Stefan-for you.

## Author Contribution

Conceptualization: DA

Study design: DA

Experimental work: DA, SA, MY, JC, LK, FC, CEOR, LJA, CGK,HAP, DJT

Molecular analyses (RNA sequencing, lipidomics): AK, MRB, SD, ZS

Bioinformatic and in silico analyses: AK, MRB, ZS, FC, SD

Data curation and statistical analysis: DA, SA, JJJJM, SMH, AK, DJT, MY,

Figure preparation and visualization: DA, ZS, AK

Manuscript drafting: DA

Manuscript review and editing: SA, AK, MY, FC, JC, CEOR, MRB, ZS, LJA, CGK, SD, HAP, JJJJM, CT, AJM, MPM, DMS, SMH, DJT, DA

Supervision: DA, DJT, CT, AJM, MPM

Funding acquisition: DA

