## Supplementary material for "Targeting immunometabolic pathways with AZD1656 alleviates inflammation and metabolic dysfunction in type 2 diabetic cardiomyopathy": Online Supplement Nov 25.pdf

| *Marker* | *Antigen* | *Fluorophore* | *Reference* | *Manufacturer* |
|---|---|---|---|---|
| Live/ Dead | Dead | 423104 | BioLegend | |
| Leukocytes | CD45 | FITC | 130-110-803 | Miltenyi |
| T cells (cytotoxic) | CD8 | Per-CP-Vio-700 | 130-128-228 | Miltenyi |
| Macrophages | F4/80 | APC | 130-131-632 | Miltenyi |
| Neutrophils | Ly6G | APC-Vio 770 | 130-128-232 | Miltenyi |
| Monocytes | LyC | VioBlue | 130-128-235 | Miltenyi |
| B cells | B220 | Vio green | 130-110-852 | Miltenyi |
| T cells (memory) | CD62L | BV785 | 564109 | BD Biosciences |
| Dendritic cells | CD11c | PE | 130-110-837 | Miltenyi |
| Monocytes/ macrophages | CD11b | PE-Vio® 615 | 130-113-811 | Miltenyi |
| T cells (regulatory) | CD4 | PE-Vio-770 | 130-127-473 | Miltenyi |
| T cells | CD3 | UV395 | 569614 | BD Biosciences |
| Treg | FoxP3 | FITC | 11-5773-82 | Invitrogen |
| Treg | CD25 | PE-Vio-770 | 130-123-028 | Miltenyi |

**Supplementary Table 1.** Antibodies used for cell staining in FACS experiments (1:200 dilution used for staining)

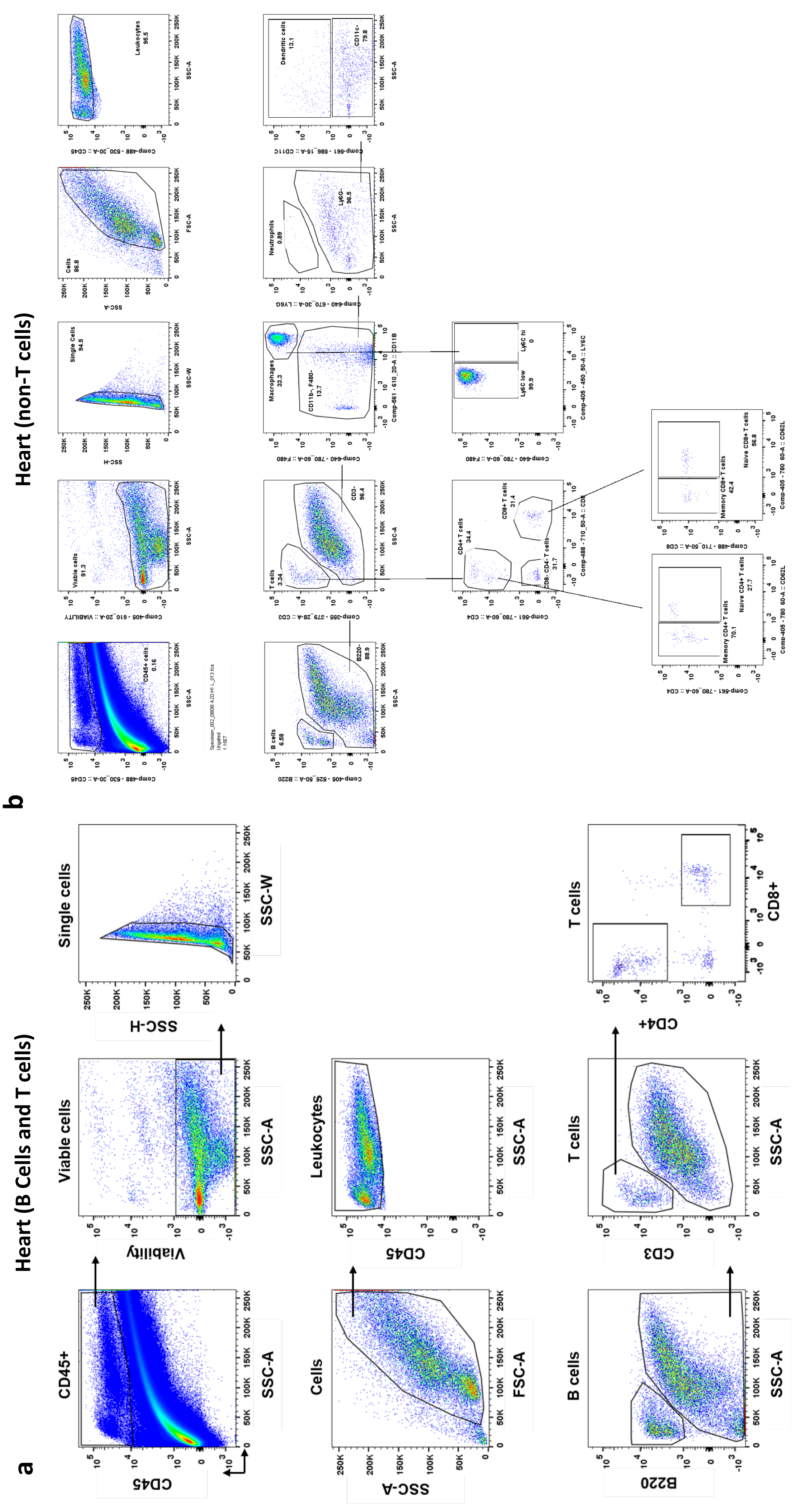


**Supplementary Figure 1.** FACS experiments representative gating strategies in a) hearts (T cell panel) b) hearts (non-T cell panel)

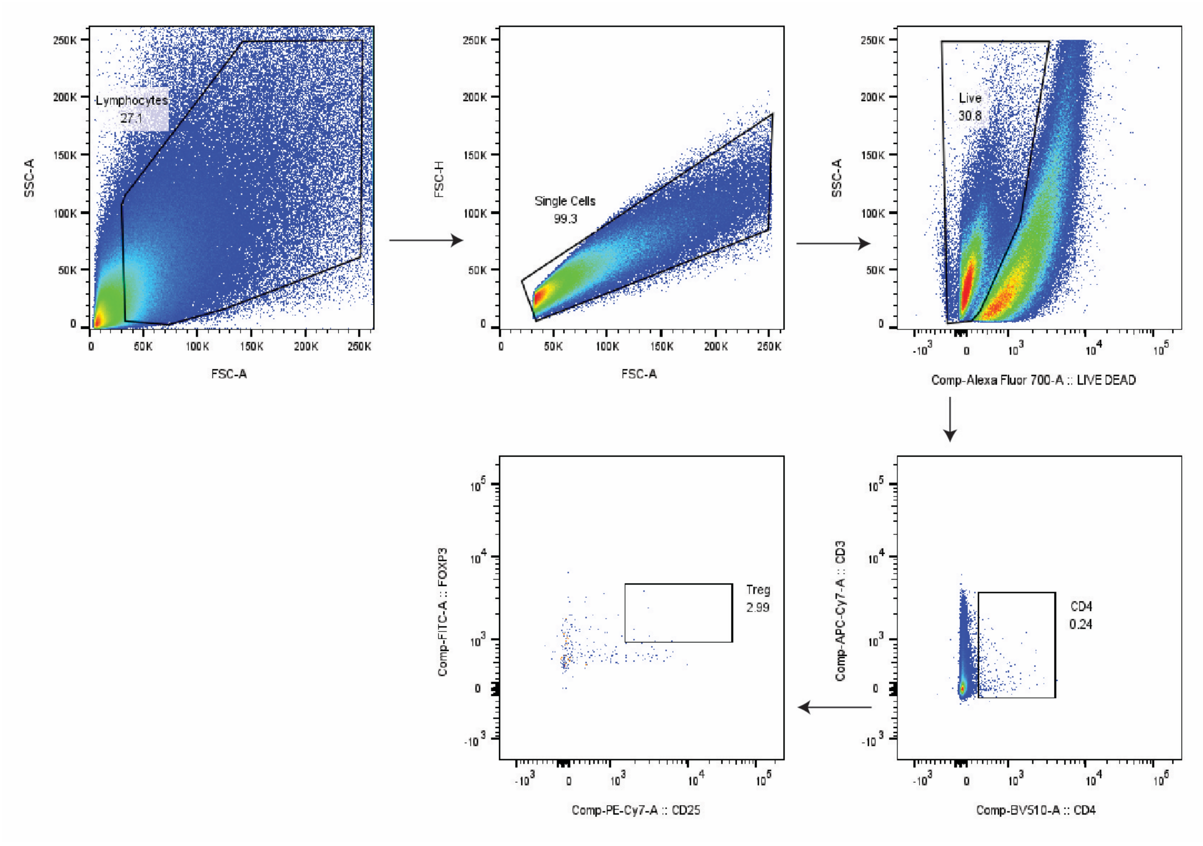


**Supplementary Figure 2.** FACS experiments representative gating strategies in hearts for detection of Tregs (CD4CD25FoxP3)

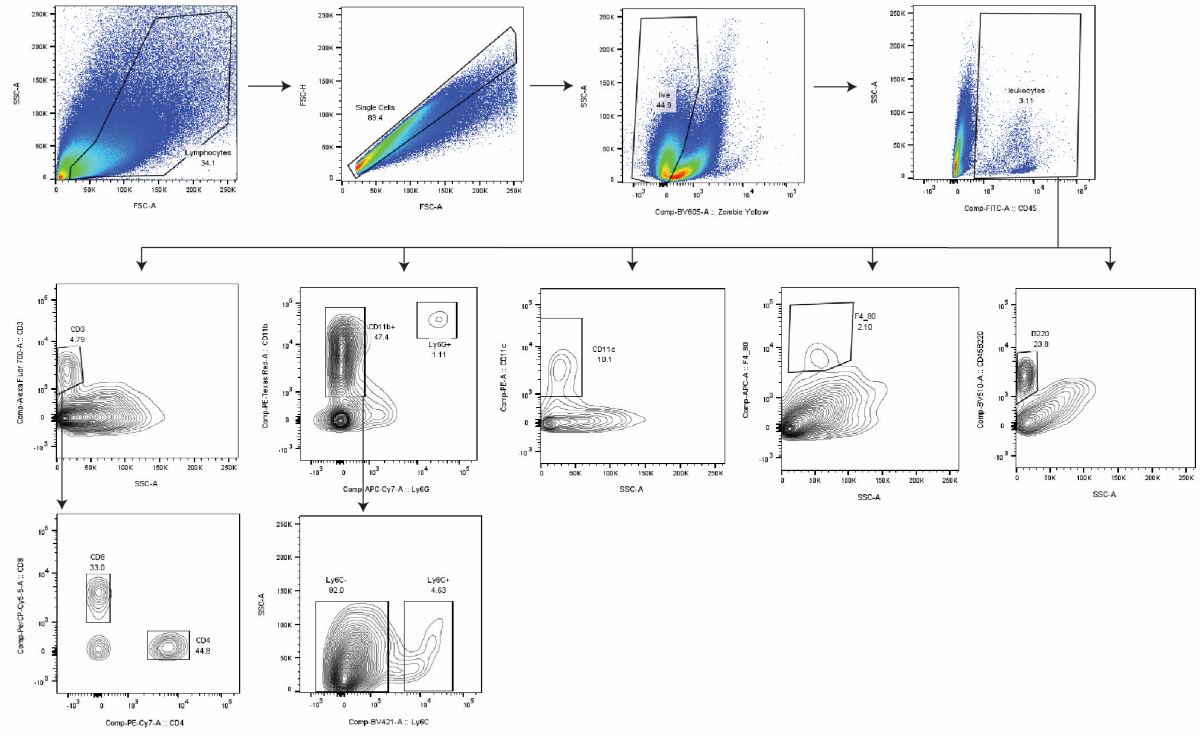


**Supplementary Figure 3. FACS experiment gating strategy for the analysis of 12 week high-fat (HFD) diet obesity model hearts**

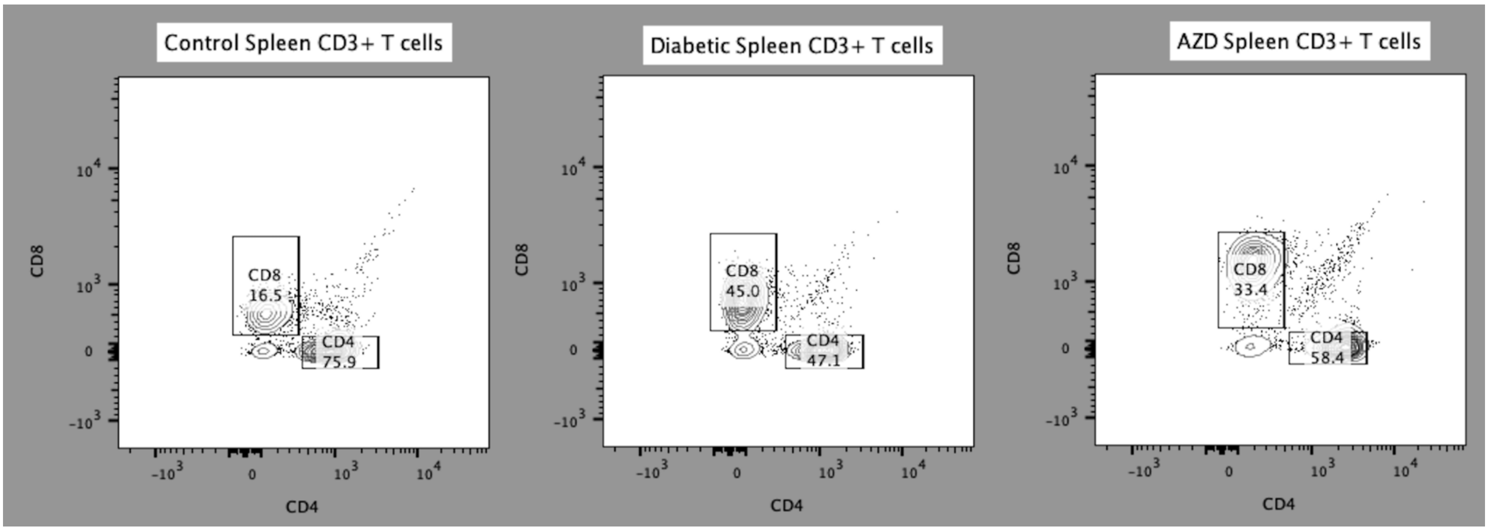


**Supplementary Figure 4. FACS experiment gating strategy for the analysis of spleen**

### Supplementary Results

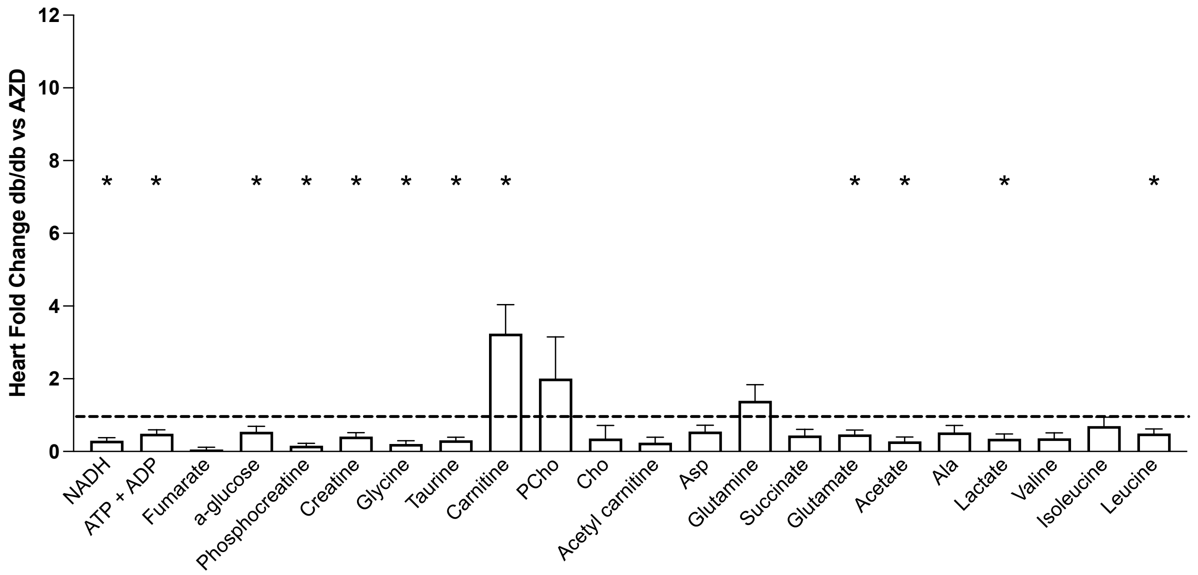


**Supplementary Figure 5. $^{1}$H NMR spectroscopy metabolomic profile of db/db vs AZD-treated db/db hearts**

Data mean+ SEM. Comparison by t test. (control n=8, db/db n=8, AZD n=5)

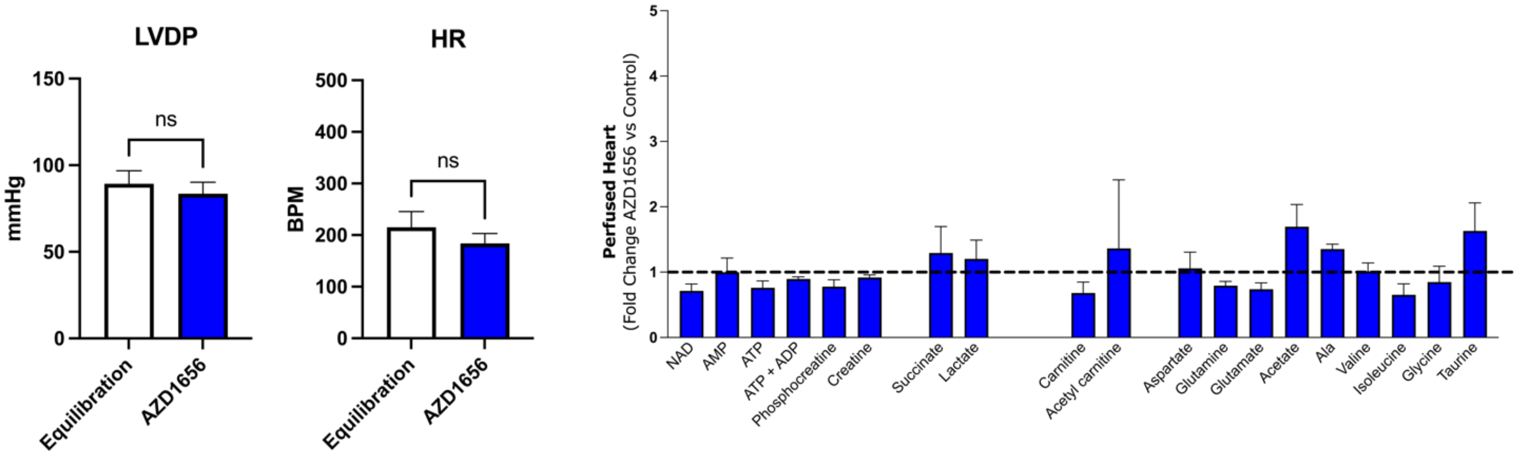


**Supplementary Figure 6. Acute AZD1656 administration has no impact on function or metabolism of hearts**

(n=4/group), data mean+ SEM. Comparison by two-tailed t test.

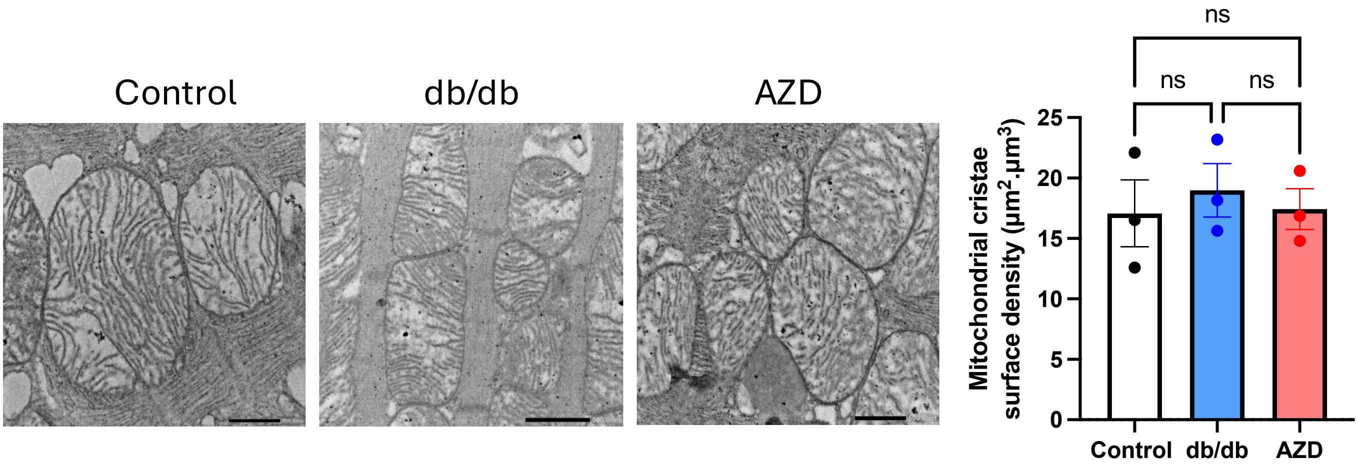


**Supplementary Figure 7. Electron microscopy assessment of mitochondrial cristae surface density** (n=3/group)

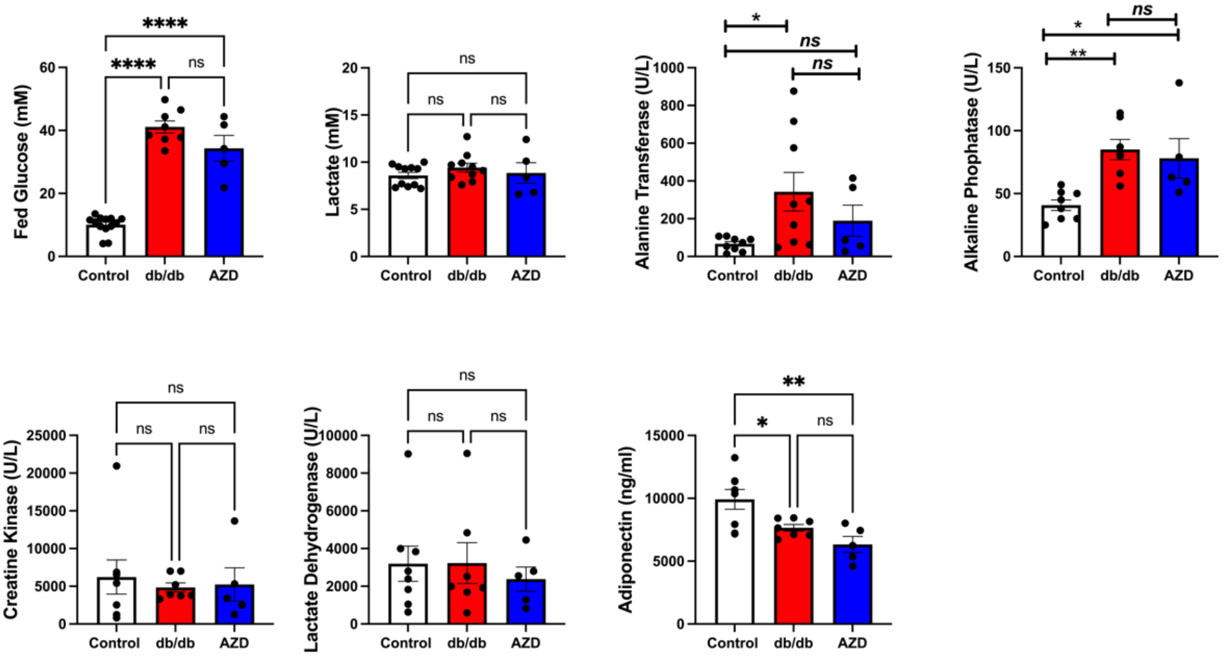


**Supplementary Figure 8. Plasma biochemical profiling**
*P<0.05, ** P<0.01, ***P<0.001 by ANOVA n=6/group

a

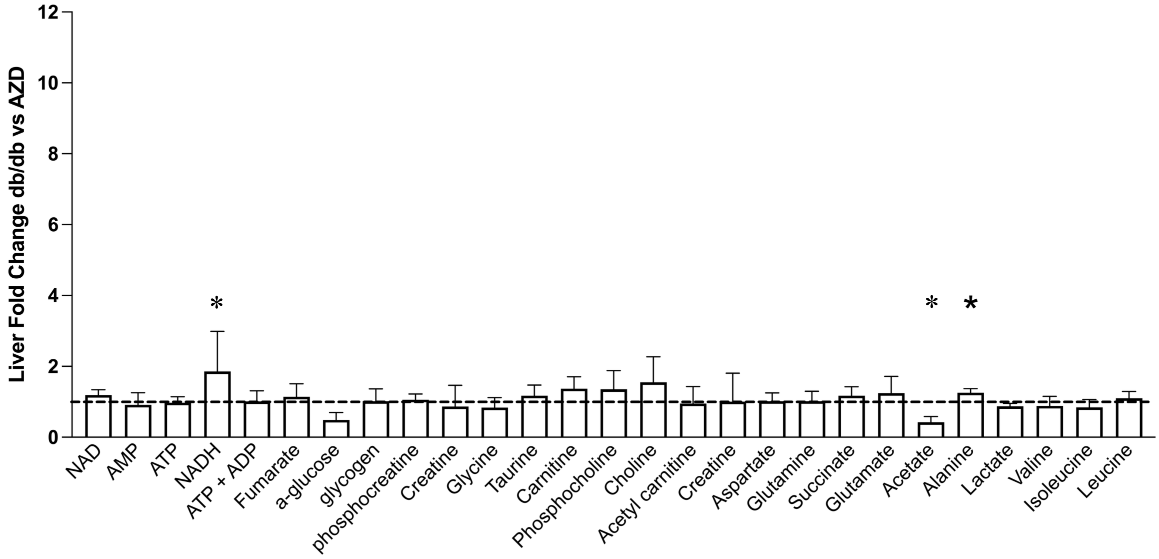


b

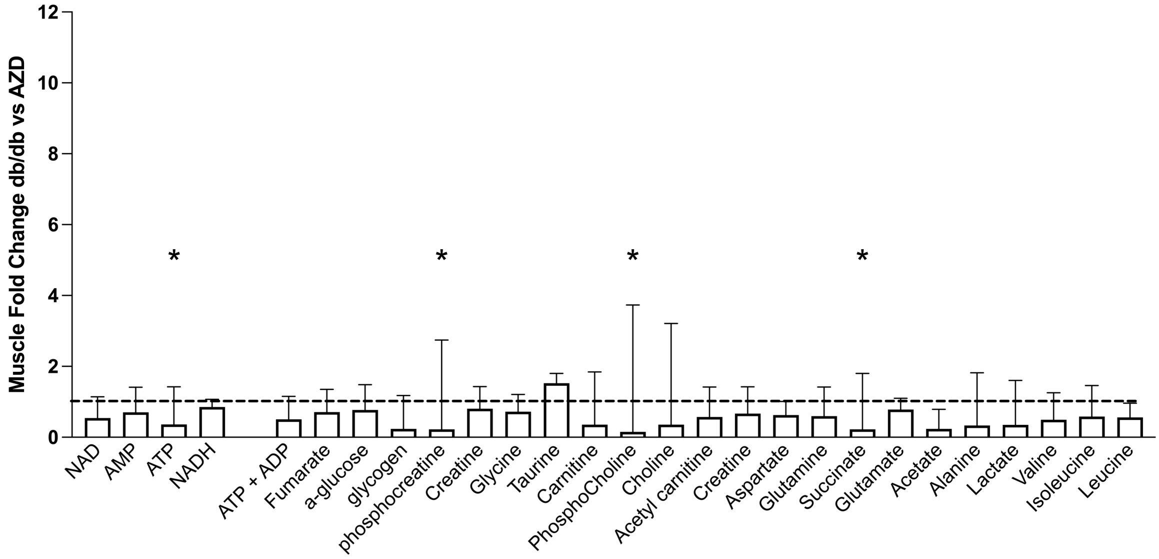


**Supplementary Figure 9. Metabolomic profile of a) liver b) muscles db/db vs AZD treated db/db animals.**

Fold change db/db vs control $^{1}$H NMR metabolomic spectroscopy profiling j) Fold change AZD vs control $^{1}$H NMR metabolomic spectroscopy profiling (Control n=6, db/db n=7, AZD n=7) Skeletal Muscle phenotype (Control n=6, db/db n=6, AZD n=5). db/db n= data mean +SEM, * P<0.05 by t-test.

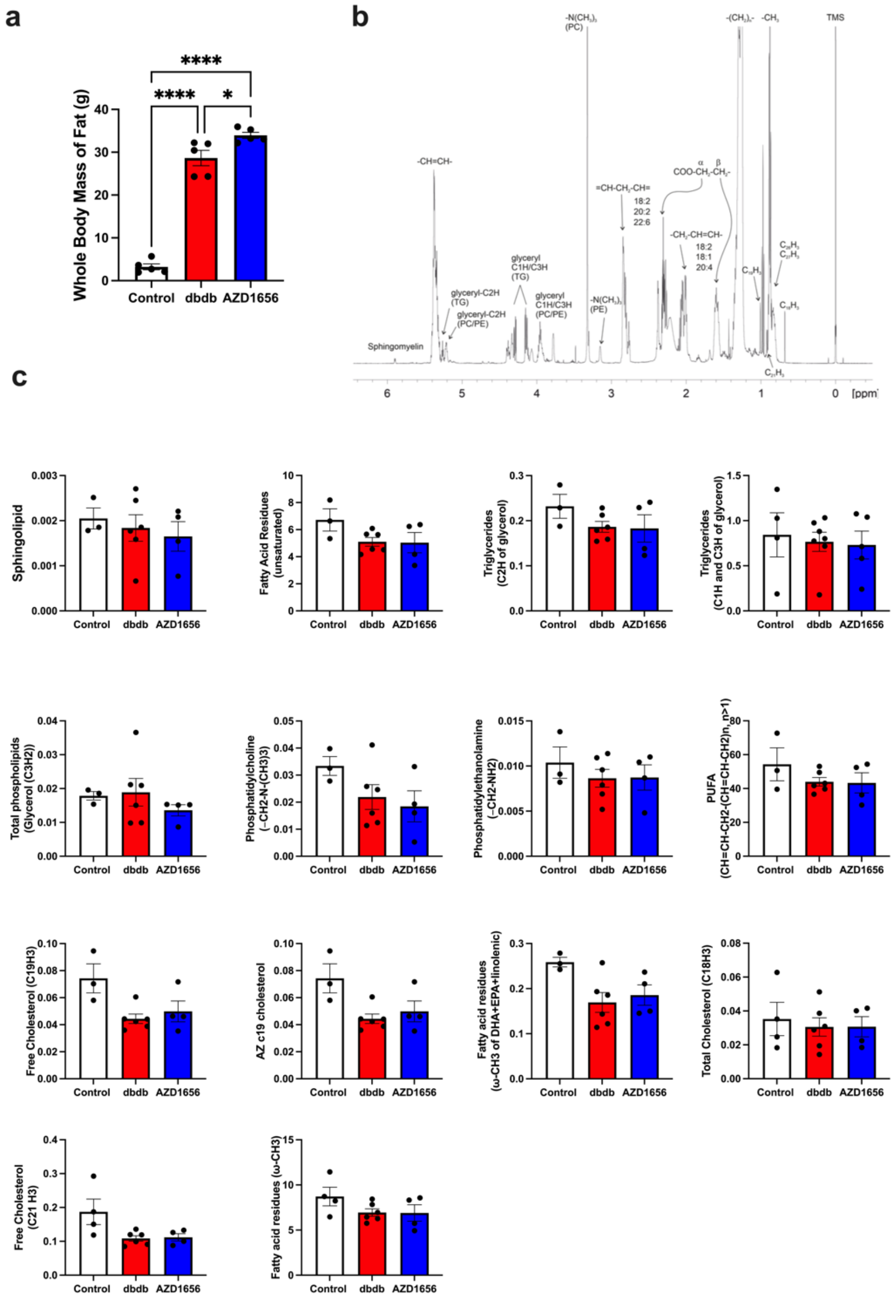

**Supplementary Figure 10. Impact of AZD1656 treatment on adipose tissue**
Whole body composition analysis b) representative 1H NMR s lipid spectrum with annotations c) adipose tissue lipid constituents quantified using $^{1}$H NMR spectroscopy (AU). n=5/group, one way ANOVA, data mean +SEM; ****p<0.0001 ****p<0.0001 *p<0.0210

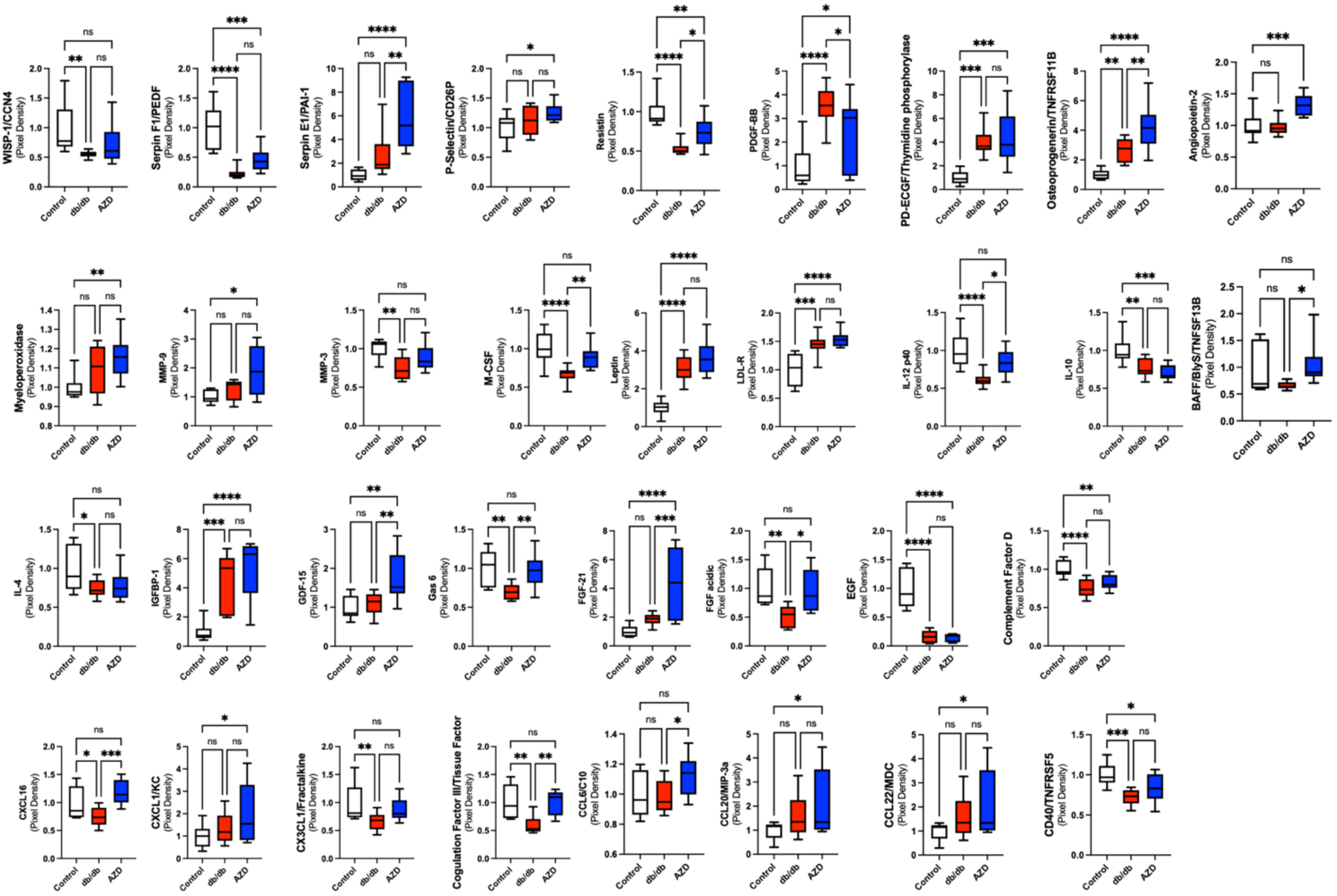


**Supplementary Figure 11. Circulating cytokine profile**
data mean+ SEM. Comparison by ANOVA n=6/group

| | Control<br>(n=6) | HFD<br>(n=4) |
|---|---|---|
| Heart rate (bpm) | 556.3 ± 26.11 | 541.6 ± 26.62 |
| Ejection fraction (%) | 67.11 ± 2.79 | 68.61 ± 2.08 |
| Fractional shortening (%) | 36.86 ± 2.15 | 38.00 ± 1.64 |
| Left ventricle internal diameter, diastole (mm) | 3.80 ± 0.19 | 3.81 ± 0.09 |
| Left ventricle internal diameter, systole (mm) | 2.43 ± 0.18 | 2.36 ± 0.11 |
| Intraventricular septum, diastole (mm) | 0.93 ± 0.05 | 0.87 ± 0.08 |
| Intraventricular septum, systole (mm) | 1.42 ± 0.08 | 1.35 ± 0.08 |
| E/A | 1.39 ± 0.03 | 1.40 ± 0.04 |
| E/E’ | 25.61 ± 1.90 | 28.64 ± 1.44 |
| Isovolumetric relaxation time (ms) | 9.20 ± 0.96 | 8.67 ± 0.81 |
| Mitral valve deceleration time (ms) | 22.98 ± 1.32 | 20.47 ± 2.13 |

**Supplementary Table 2. Echocardiography of control and 12 week high fat diet feeding-induced obesity**

Analysed by Student’s t-test, *p<0.05 vs. Sham. Data displayed as mean ± SEM

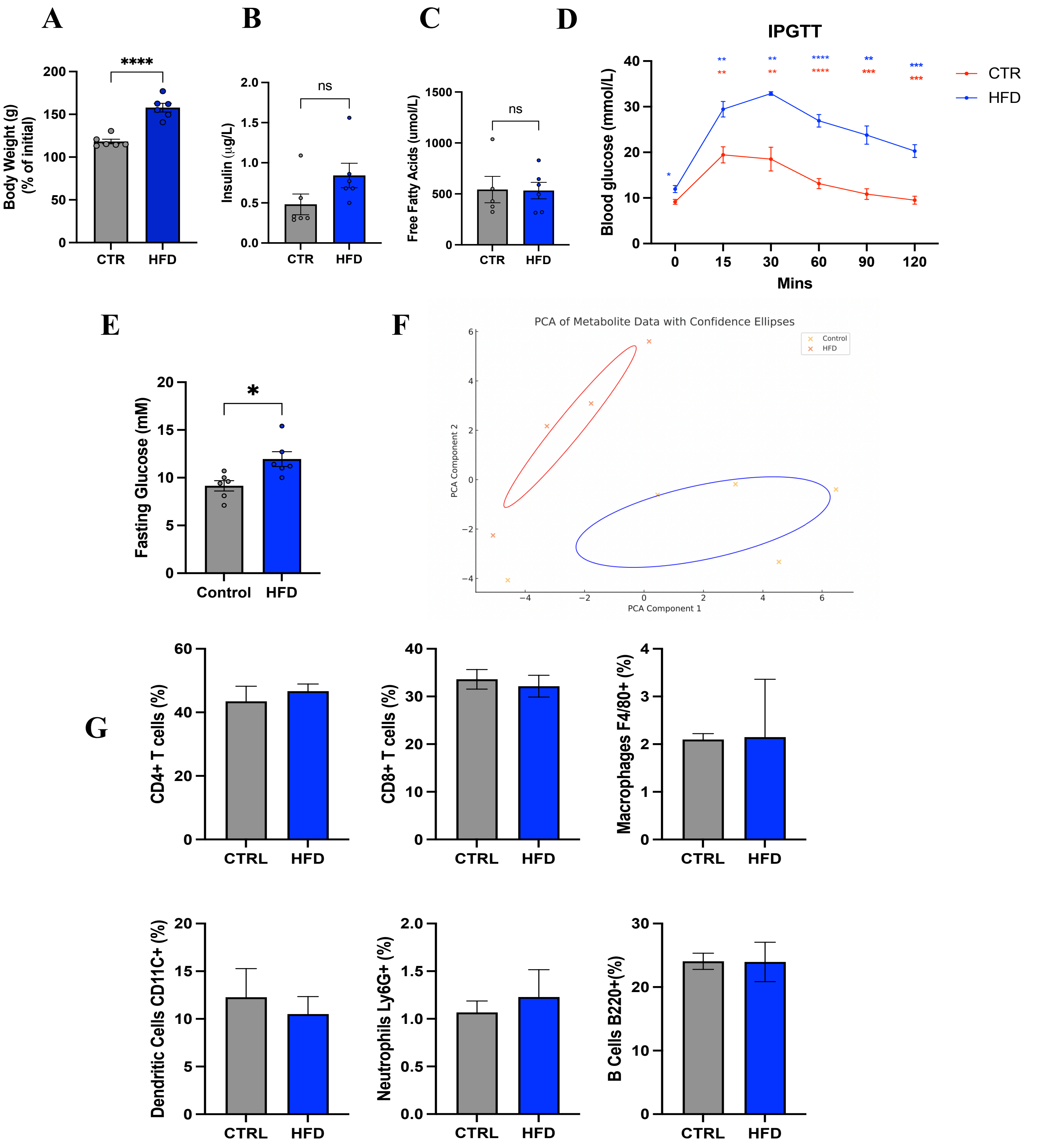


**Supplementary Figure 12. Physiological profile of 12 week HFD-fed model of pre-diabetes A)**End point body weight **B)** Endpoint plasma insulin, P=0.054 **C)** Endpoint Plasma free fatty acids **D)** IPGTT **E)** Fasting plasma glucose **F)** PCA plot of cardiac metabolomic profile analysed by LC/MS **G)** Myocardial inflammatory cell profile assessed by flow cytometry n=6/group, male, C57/BL6 mice, age at endpoint 20 weeks. Flow experiment n=3/group IPGTT- intraperitoneal glucose tolerance test. *P<0.05 ***P<0.001 ****P<0.0001 by t -test or ANOVA (D)

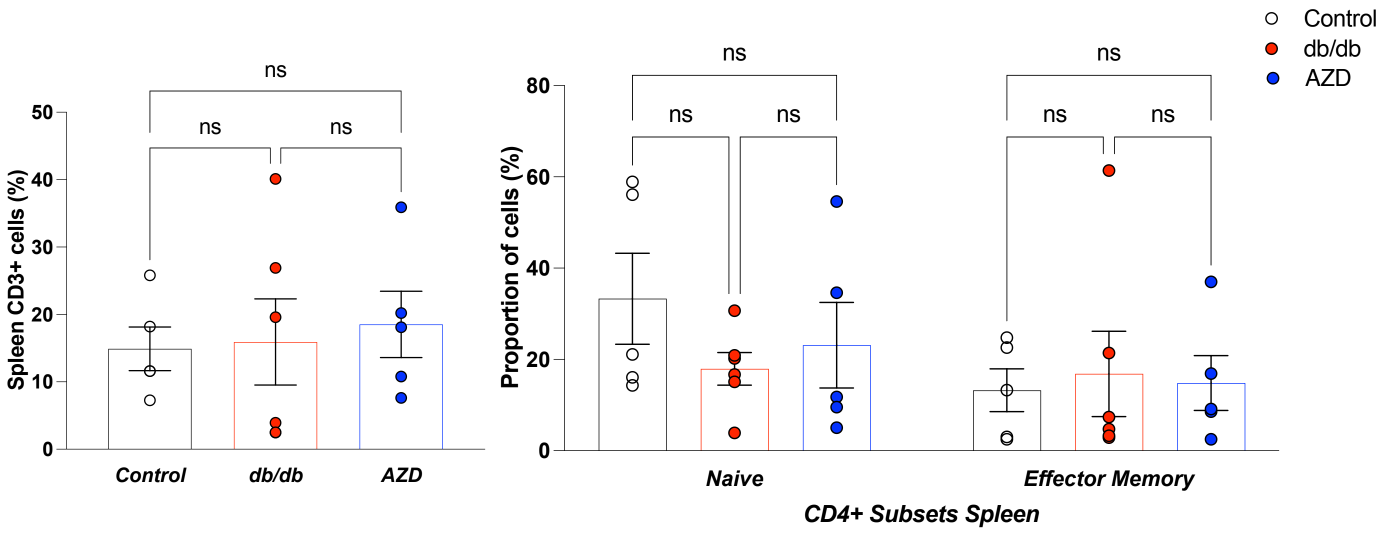


**Supplementary Figure 13. FACS assessment of the spleen T cell profile**

(Control n=5, db/db n=6 AZD1656 n=5)

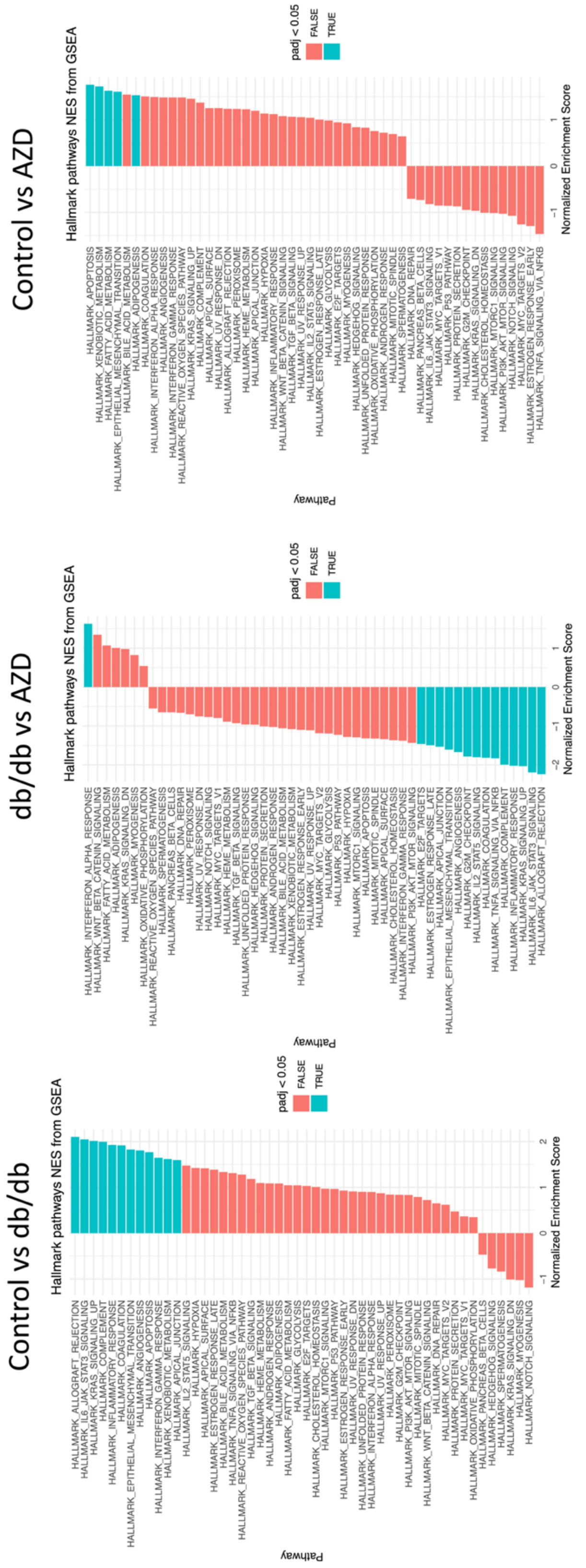


**Supplementary Figure 14. G profiler analysis of the cellular pathways identified from RNA cardiac sequencing data (DEG gene expression)**

n=6/group. Raw RNA seq expression data provided in Supplementary Data Table 1.

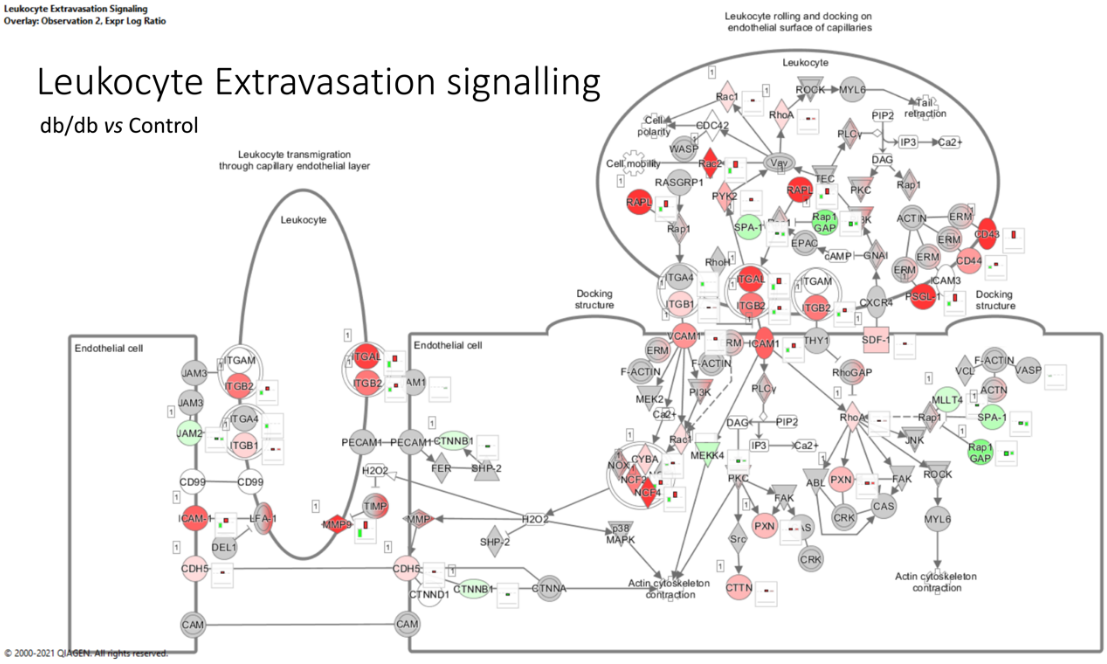


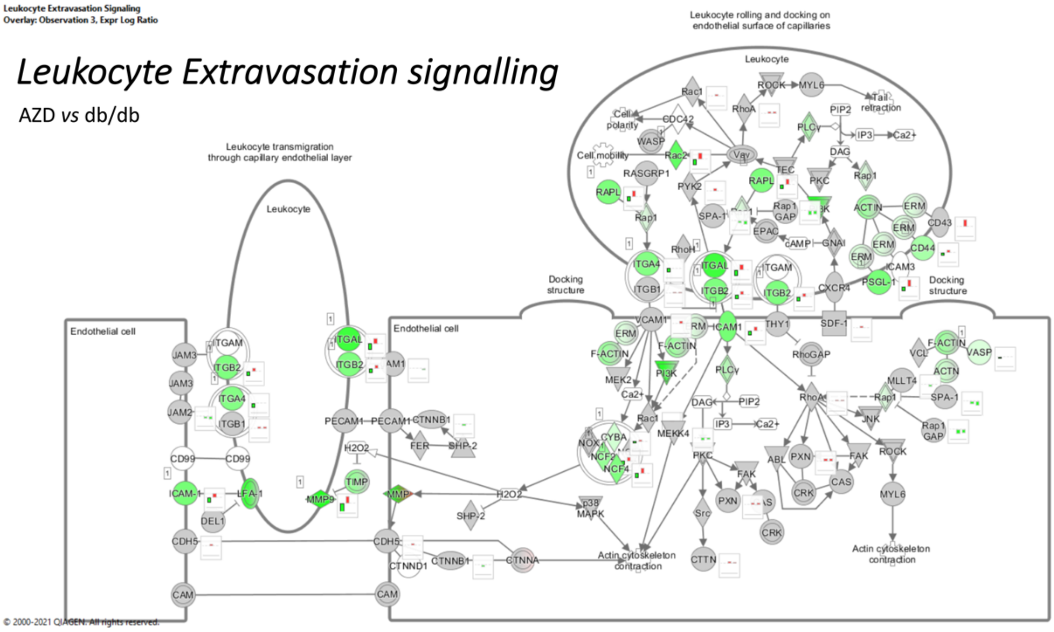


**Supplementary Figure 15. AZD treatment normalizes aberrant myocardial leukocyte extravasation signalling pathway**

Qiagen Ingenuity Pathway Analysis (IPA) based on RNA sequencing results (n=6/group). IPA utilise hypergeometric test and Benjamini-Hochberg p-value correction to identify all ontology and pathway terms that contain a statistically greater number of genes in common with an input list than expected by chance. Red indicates altered expression and pathway impairment. Green signifies gene expression normalisation.

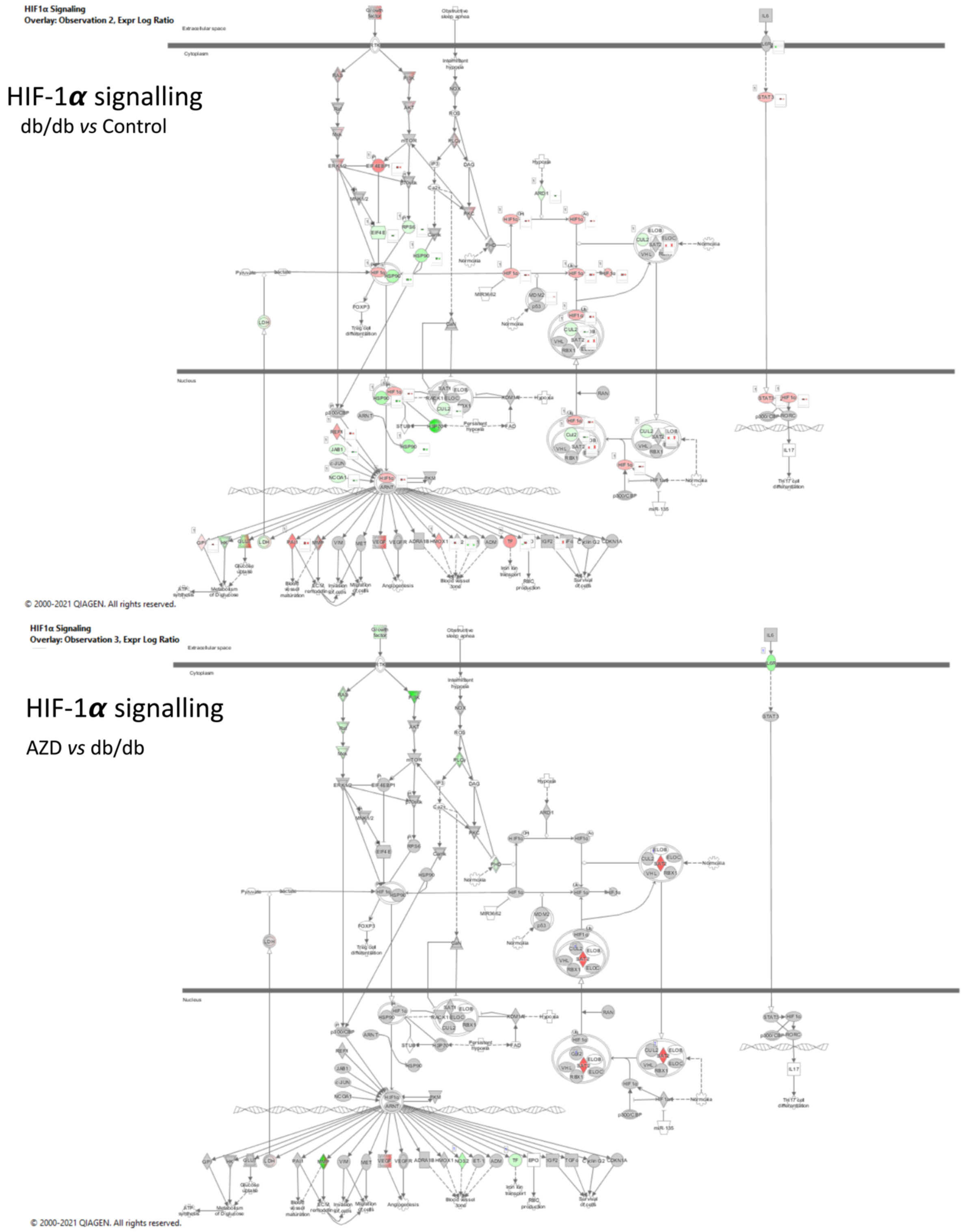


**Figure 16. AZD treatment normalizes myocardial HIF-1α signalling pathway in dbCM**
Qiagen Ingenuity Pathway Analysis (IPA) based on RNA sequencing results (n=6/group). IPA utilise hypergeometric test and Benjamini-Hochberg p-value correction to identify all ontology and pathway terms that contain a statistically greater number of genes in common with an input list than expected by chance.

*Nrf 2-mediated Oxidative Stress Response*

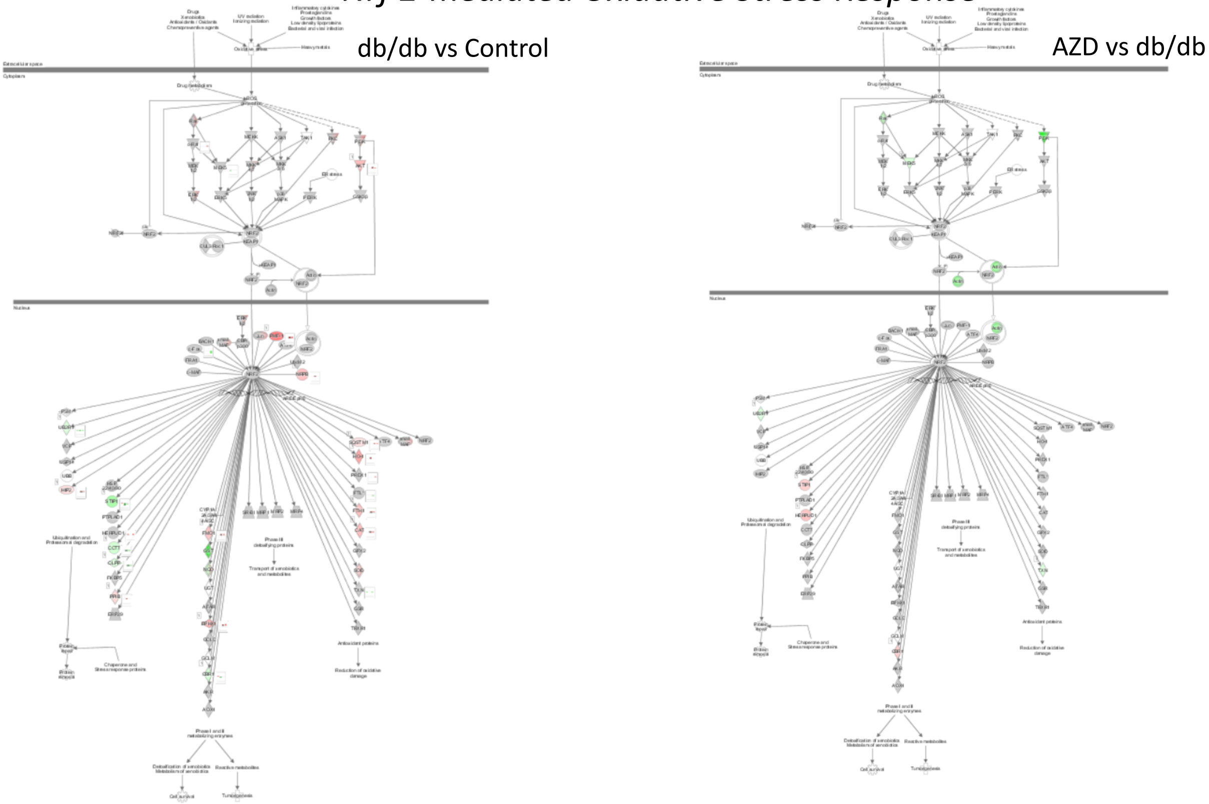


**Figure 17. AZD treatment normalizes myocardial Nrf2 signalling pathway in dbCM**
Qiagen Ingenuity Pathway Analysis (IPA) based on RNA sequencing results (n=6/group). IPA utilise hypergeometric test and Benjamini-Hochberg p-value correction to identify all ontology and pathway terms that contain a statistically greater number of genes in common with an input list than expected by chance.
